# Large-scale mapping and systematic mutagenesis of human transcriptional effector domains

**DOI:** 10.1101/2022.08.26.505496

**Authors:** Nicole DelRosso, Josh Tycko, Peter Suzuki, Cecelia Andrews, Aradhana, Adi Mukund, Ivan Liongson, Connor Ludwig, Kaitlyn Spees, Polly Fordyce, Michael C. Bassik, Lacramioara Bintu

**Affiliations:** Biophysics Program, Stanford University, Stanford, CA 94305, USA; Department of Genetics, Stanford University, Stanford, CA 94305, USA; Department of Bioengineering, Stanford University, Stanford, CA 94305, USA; Department of Developmental Biology, Stanford University, Stanford, CA 94305, USA; Department of Biology, Stanford University, Stanford, CA 94305, USA; ChEM-H Institute, Stanford University, Stanford, CA 94305, USA; Chan Zuckerberg Biohub, San Francisco, CA 94110, USA

## Abstract

Human gene expression is regulated by over two thousand transcription factors and chromatin regulators^1,2^. Effector domains within these proteins can activate or repress transcription. However, for many of these regulators we do not know what type of transcriptional effector domains they contain, their location in the protein, their activation and repression strengths, and the amino acids that are necessary for their functions. Here, we systematically measure the transcriptional effector activity of >100,000 protein fragments (each 80 amino acids long) tiling across most chromatin regulators and transcription factors in human cells (2,047 proteins). By testing the effect they have when recruited at reporter genes, we annotate 307 new activation domains and 592 new repression domains, a ∼5-fold increase over the number of previously annotated effectors^3,4^. Complementary rational mutagenesis and deletion scans across all the effector domains reveal aromatic and/or leucine residues interspersed with acidic, proline, serine, and/or glutamine residues are necessary for activation domain activity. Additionally, the majority of repression domain sequences contain either sites for SUMOylation, short interaction motifs for recruiting co-repressors, or are structured binding domains for recruiting other repressive proteins. Surprisingly, we discover bifunctional domains that can both activate and repress and can dynamically split a cell population into high- and low-expression subpopulations. Our systematic annotation and characterization of transcriptional effector domains provides a rich resource for understanding the function of human transcription factors and chromatin regulators, engineering compact tools for controlling gene expression, and refining predictive computational models of effector domain function.

## Introduction

Human gene expression is regulated by a constellation of over two thousand proteins known as transcription factors (TFs) and chromatin regulators (CRs). TFs bind genomic DNA site-specifically^1^, and CRs recognize DNA and histone modifications^5^. Both classes of proteins contain effector domains that recruit other macromolecules to activate or repress transcription. Consequently, mapping the location of effector domains within these thousands of proteins is an essential resource for decoding the functional properties of the human genome. Large scale efforts have mapped where in the genome TFs and CRs bind^6,7^. However, equivalent maps of human transcriptional effector domains are incomplete: we are currently missing effector domain annotations for about 60% of the human TFs^8^.

Moreover, the sequence characteristics of what makes a good human activation or repression domain are still under current investigation. Activation domains (ADs) are often disordered and are typically characterized by their amino acid compositions^9,10^. Most AD sequence characteristics have been elucidated from yeast, where all ADs consist of a mix of acidic and hydrophobic residues^11–13^. The acidic residues are thought to keep the hydrophobic residues exposed to contact co-activators, an idea known as the acidic exposure model^14,15^. In addition to acidic activators, some human ADs have non-acidic compositional biases, such as glutamine-, proline-, serine-, glycine-, and alanine-rich sequences^16–19^. It remains unclear how many non-acidic human ADs there are and how they work.

Repression domains (RDs), on the other hand, are less disordered^8^. As a result, RDs are not typically described by their sequence compositions. Instead, a more useful description of RDs has been categorization of their structural domains and repression mechanisms. For example, the family of hundreds of KRAB domains generally recruit the scaffold protein KAP1, which represses transcription and creates H3K9me3-associated heterochromatin by further recruitment of histone deacetylases, histone methylase SETDB1, and heterochromatin protein HP1^20,21^. Systematic categorization of human RD sequences remains incomplete.

One useful assay for characterizing individual protein effector domains and mutants that test specific sequence requirements consists of recruitment of domains to reporter genes (reviewed in^8^). This approach has been extended from recruiting single domains to high-throughput assays in yeast^11,12,14,22^, drosophila^23–25^, and human cells with a subset of transcriptional domains^4^ or a subset of full length transcription factors^26^. These works have extended our list of effector domains and have set the stage for systematically mapping the effector domains across the thousands of human transcriptional proteins.

In order to map the human effector domains at unprecedented scale and resolution, here, we use a high-throughput reporter assay to measure the transcriptional activity of 113,528 protein fragments tiling across 2,047 chromatin regulators and transcription factors, the largest high-throughput assay for protein function performed in human cells to date. Using rational mutagenesis and deletion scanning at scale, we elucidate necessary sequence properties for both activation and repression domains. We find that AD sequences that are glutamine-, proline-, and serine-rich behave similarly to acidic sequences: the glutamines, prolines, and serines that are necessary for activity are the ones that are interspersed with hydrophobic residues. Additionally, our data suggest the pervasive role that SUMOylation and zinc finger domains play in causing repression for hundreds of RDs. Finally, we uncover a new set of bifunctional domains, some of which are capable of simultaneously enhancing and silencing expression from a single promoter.

## High-throughput mapping of effector domains

To map the effector domain locations in human TFs and CRs, we selected 1,292 human TFs from the Lambert 2018 dataset^1^, 735 CRs from the EpiFactors Database^2^, and added 20 genes with GO terms matching chromatin and histone regulation (**Supplementary Table 1, Methods**). To make this library’s size feasible for high throughput measurements, we excluded 473 proteins that we have previously characterized with HT-recruit^4^: a set of 129 CRs^4^ and 344 KRAB-containing TFs. For each TF and CR in our list, we synthesized DNA sequences encoding 80 amino acid (aa) segments that tile across the full-length protein (hereafter CRTF tiling library) with a 10 aa step size between segments (**Fig. 1a**). In addition, we included 2,000 random 80 aa protein sequences as negative controls, 10 previously validated effector domains^4^ as positive controls, and a deletion scan across 50 UniProt annotated ADs as a pilot test of sequence perturbations (**Methods**).

**Fig 1.**
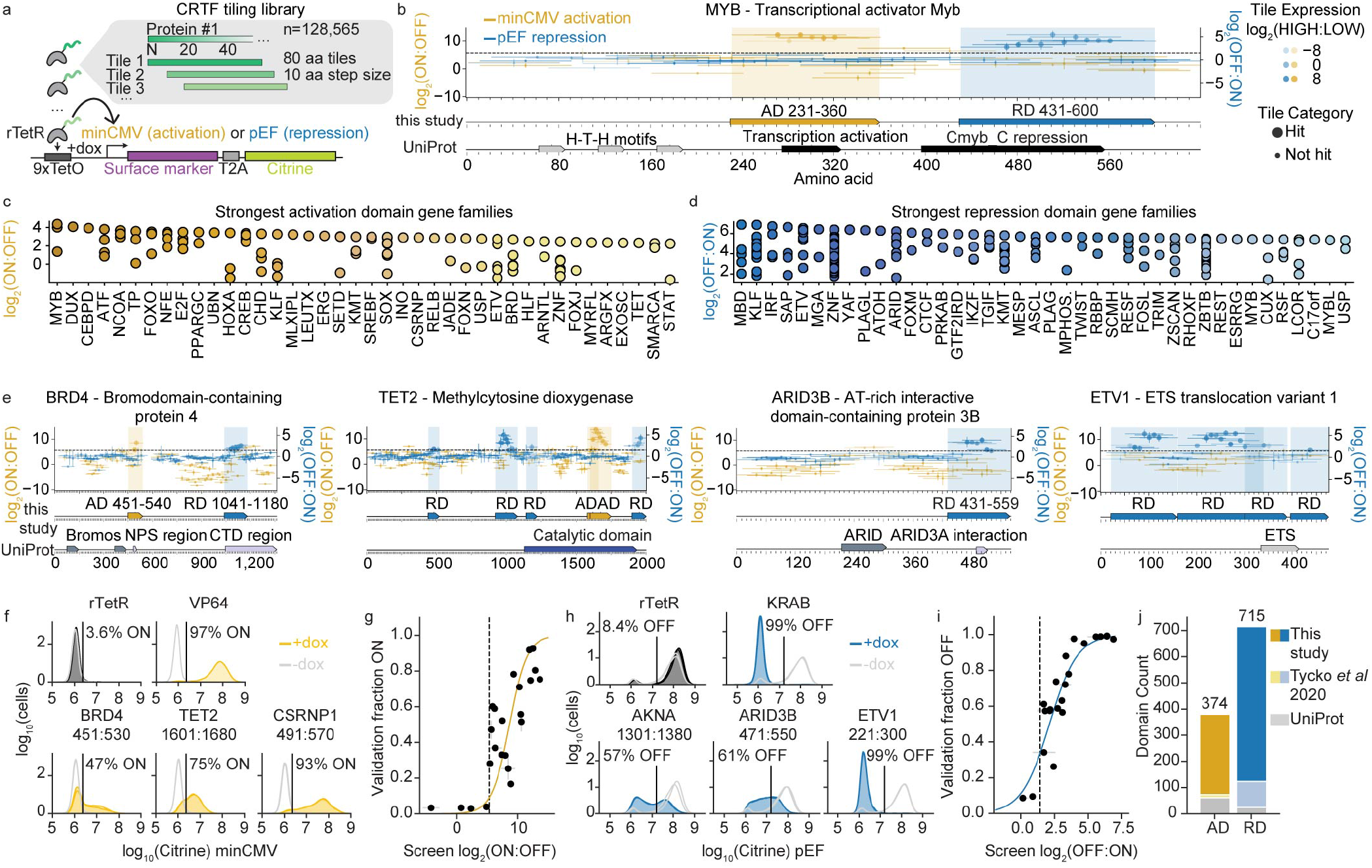
High-throughput tiling screen across 2,047 human TFs and CRs finds hundreds of undiscovered effector domains. **a**, Each protein’s activating and repressing regions are identified by partitioning the full-length sequence into 80 aa tiles. Each tile is fused to rTetR-3xFLAG, and the library is delivered with lentivirus to K562 cells containing a magnetic-fluorescent reporter (surface marker and citrine) stably integrated in the safe harbor *AAVS1* locus. The reporter contains 9 TetO binding sites for recruiting a rTetR-tile fusion upon dox addition (+dox). Activation is measured after recruitment upstream of a minimal promoter (minCMV) for 2 days, and repression after recruitment upstream of a constitutive promoter (pEF) for 5 days. **b**, Activation (at minCMV, yellow) and repression (at pEF, blue) tiling enrichment scores overlap previously annotated MYB effector domains (bottom, black, sourced from UniProt). Each horizontal yellow and blue line represents an 80 aa tile, and each vertical error bar is the standard error from two biological replicates. The dashed horizontal line represents the hit calling threshold based on random controls (Methods). Points with larger marker sizes were hits in the second (validation) screen. FLAG-stained expression levels are plotted as the hue, with higher expressing-tiles in darker hues. Effector domains identified in this study are annotated as contiguous regions at the bottom: yellow bars for ADs and blue bars for RDs (Methods). **c-d**, Distribution of the strongest effector domains from the top 40 gene families. Enrichment scores are from the validation screen (Extended Data Fig. 3), measured for the maximum activating/repressing tile within each domain. **e**, Tiling results for BRD4, TET2, ARID3B, and ETV1. **f**, Individual validations of activating tiles after 2 days of recruitment (+dox). Untreated cells (gray) and dox-treated cells (colors) shown with two biological replicates in each condition. Vertical line is the citrine gate used to determine the fraction of cells ON (written above each distribution). rTetR alone is a negative control and VP64 is a positive control. **g**, Comparison between individually recruited and screen measurements with logistic model fit plotted as solid line (r^2^=0.67, N=23). Error bars are the standard error for 2 biological replicates (screens and validations) and dashed line is the hits threshold. **h**, Individual validations of repressing tiles after 5 days of recruitment (n=2). KRAB is a positive control. **i**, Comparison between individually recruited and pEF promoter screen measurements (r^2^=0.84, N=22). **j**, Domain counts that are new (dark gold and blue), overlap UniProt annotations (gray), or overlap prior HT-recruit screen results^4^ (light gold and blue). Total is shown above each bar. RDs are annotated from tiles that were hits in both pEF and PGK promoter screens (Extended Data Fig. 4).

This library, consisting of 128,565 sequences, was cloned into a lentiviral vector, where each protein tile is expressed as a fusion protein with rTetR (a doxycycline inducible DNA binding domain), and delivered to K562 cells containing a reporter with binding sites for rTetR^4^ (**Fig. 1a, Methods**). The reporter gene is driven by either a minimally active minCMV promoter for identifying activators, or constitutively active pEF promoter for finding repressors. To simultaneously measure the transcriptional effector function of these sequences, we used a high-throughput recruitment assay we recently developed: HT-recruit^4^. Briefly, the library was cloned and delivered as a pool at a low lentiviral infection rate, such that each cell contains a single rTetR-tile. After treating the cells with doxycycline, which recruits each CRTF tiling library member to the reporter, we magnetically separated the cells into ON and OFF populations, extracted genomic DNA, and sequenced the tiles to identify sequences that were enriched in the ON or OFF cell populations (**Extended Data Fig. 1a-b**). Activating tiles are enriched in the ON population, while repressing tiles are enriched in the OFF population. Each screen was reproducible across two biological replicates (**Extended Data Fig. 1c-d**). Using the random negative controls, we drew thresholds for calling hits (**Extended Data Fig. 1c-d, Methods**). 90% and 92% of the positive control domains for activation and repression^4^, respectively, were hits above this threshold, as expected (**Supplementary Table 1**). We identified an additional subset of shared tiles (n=175) that were only hits in this repression screen (**Extended Data Fig. 1e**) and whose activity validated in individual flow cytometry experiments (**Extended Data Fig. 1f**). Overall, these results demonstrated HT-recruit reliably identified transcriptional effectors while using an order-of-magnitude larger library than our previous experiments^4^.

As measured transcriptional strength depends not only on the intrinsic potential of the sequence but also on the levels at which individual tiles are expressed, we measured expression of all protein tiles. All our library members contain a 3xFLAG tag, allowing us to measure each fusion protein’s expression levels by staining with an anti-FLAG antibody, FACS sorting the cells into FLAG HIGH and LOW populations (**Extended Data Fig. 2a**), and measuring the abundance of each member in the two populations by sequencing the domains (**Extended Data Fig. 2b**). Our FLAG screen scores correlate well with individual validations (**Extended Data Fig. 2c**). These expression measurements were used when annotating effector domains, for example allowing us to identify and filter out false negative library members that have lower activation or repression scores due to low expression (**Extended Data Fig. 2d, Methods**).

**Fig. 2.**
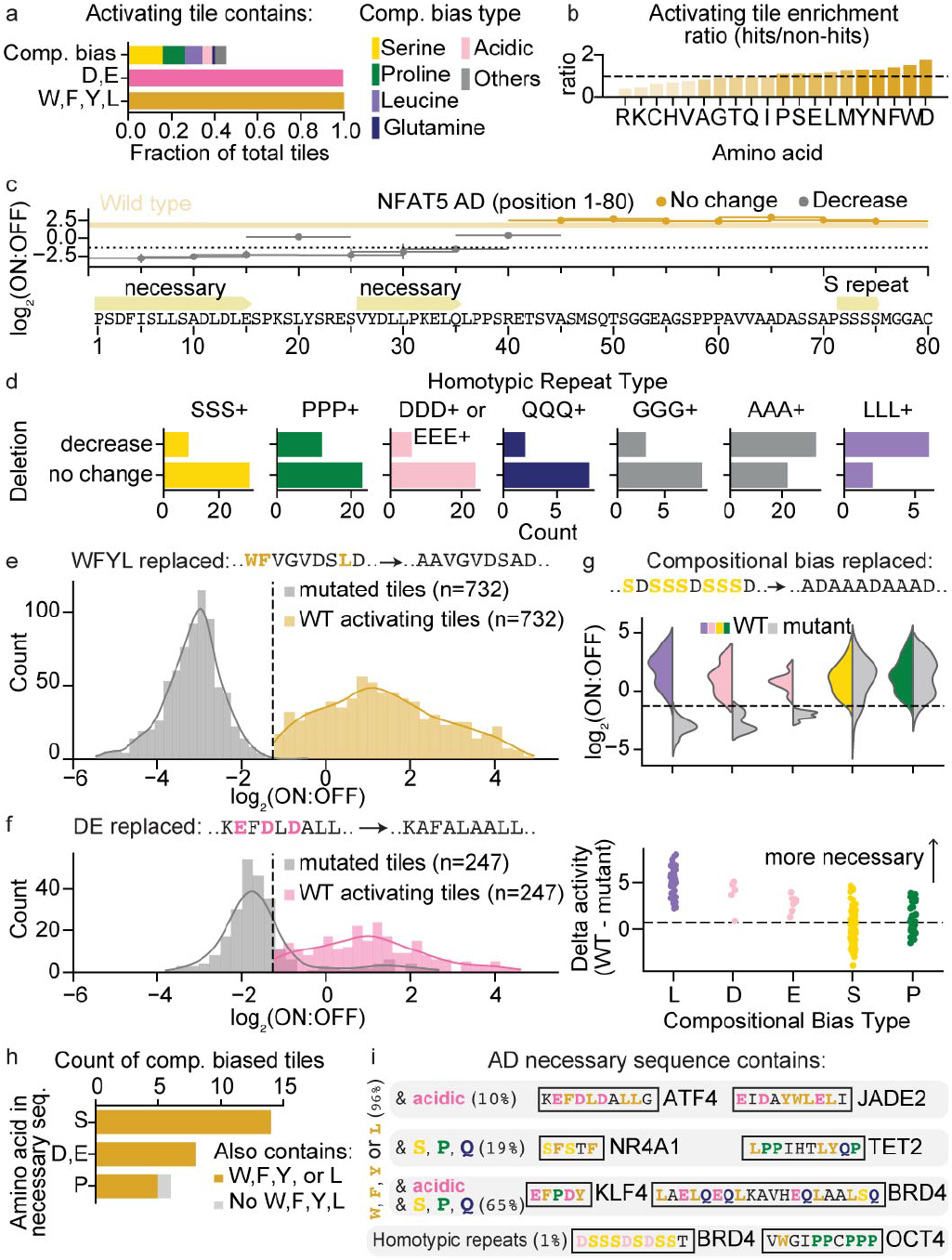
Hydrophobic amino acids that are interspersed with acidic, serine, proline or glutamine residues are necessary for AD activity. **a**, Fraction of activating tiles that contain a compositional bias (amino acids that appeared at least 12 times in the 80 aa, i.e. 15% of the sequence). Number of tiles in each compositional bias type: serine=132, proline=85, leucine=69, acidic=36, glutamine=10, others are alanine=21, glycine=13, asparagine=6, and methionine=3. **b**, Enrichment ratio for each amino acid across all activating tile sequences. Ratios were computed by counting the abundance of each amino acid in the hit sequences, and normalizing by the length and total number of sequences. Randomly sampled 10,000 non-hit 80 aa sequences were similarly calculated and the enrichment ratio was calculated by dividing the hits by non-hits. Horizontal dashed line is at a ratio of 1. **c**, Deletion scan across NFAT5’s AD. Yellow rectangle represents WT enrichment score, with the standard error between two biological replicates represented as the height. If the deletion’s score is lower than 2 times the average standard error for measuring a deletion, it’s binned as “decrease.” Otherwise it’s binned as “no change.” **d**, Counts of deletion sequences containing a homotypic repeat of 3 or more amino acids of the indicated type binned according to their effect on activity compared to the WT sequence: decrease or no change upon deletion. Probability we would observe serine ratio p=7.58e-3, proline=5.17e-2, acidic=6.57e-3, glutamine=1.73e-3, glycine=6.48e-2, alanine=0.699 (Fisher’s exact test compared with LLL+ distribution, two-sided) **e**, Distribution of average activation enrichment scores (2 biological screen replicates) for WT (yellow) and W,F,Y,L mutant tiles (gray) for all well-expressed W,F,Y,L-containing activating tiles. Dashed line represents the hit threshold. **f**, Distribution of average activation enrichment scores (2 biological screen replicates) for WT (yellow) and D,E mutant tiles (gray) for all well-expressed D,E-containing activating tiles. **g**, (Top) Distributions of average activation enrichment scores (2 biological screen replicates) for WT (colors) and compositional bias mutants (gray). Dashed line represents the hit threshold. (Bottom) Mutant enrichment scores subtracted from WT enrichment scores plotted for each compositional bias that was replaced with alanine. Dashed line drawn 2 times the average standard error (across all mutants) above 0. **h**, Count of all compositionally biased tiles that lost activity upon mutation that contain the compositionally biased amino acid in at least one of its necessary sequences and whether an aromatic/leucine was also present (yellow). Probability we would observe this for serine: p=3.75e-4, acidic residues: p=3.0e-3, proline residues: p=5.45e-1 (Fisher’s exact test comparing counts of tiles that had W,F,Y, or L present with a size-matched, randomly selected distribution of sequences that had no change upon deletion, two-sided). Deletion scans were only performed on the max activating tile from each AD, so only the max tile from a compositionally biased AD has a corresponding necessary sequence. **i**, Summary of findings: AD sequences that are necessary for function consist of hydrophobic amino acids that are interspersed with acidic, prolines, serines and/or glutamine residues.

To further confirm all the hits and help remove false positives, we screened a smaller library containing only the activating and repressive hit tiles (referred to as the validation screen, **Supplementary Table 1, Methods**). Because of their small size (1,055 activating tiles and 7,939 repressive tiles), these screens had better magnetic separation purity (**Extended Data Fig. 3a-b**), and the libraries could be screened at 10-fold higher coverage, which resulted in higher reproducibility than the original, larger screens (**Extended Data Fig. 3c-d**), and even better correlation between screen scores and individual validations (**Extended Data Fig. 3e-f**). Encouragingly, about 80% of the original hits also were confirmed as hits in these validation screens (**Supplementary Table 1, Extended Data Fig. 3c-d**). We only considered these confirmed sequences in subsequent analyses: 830 activation and 6,755 repression tiles (**Supplementary Table 1**).

**Fig. 3.**
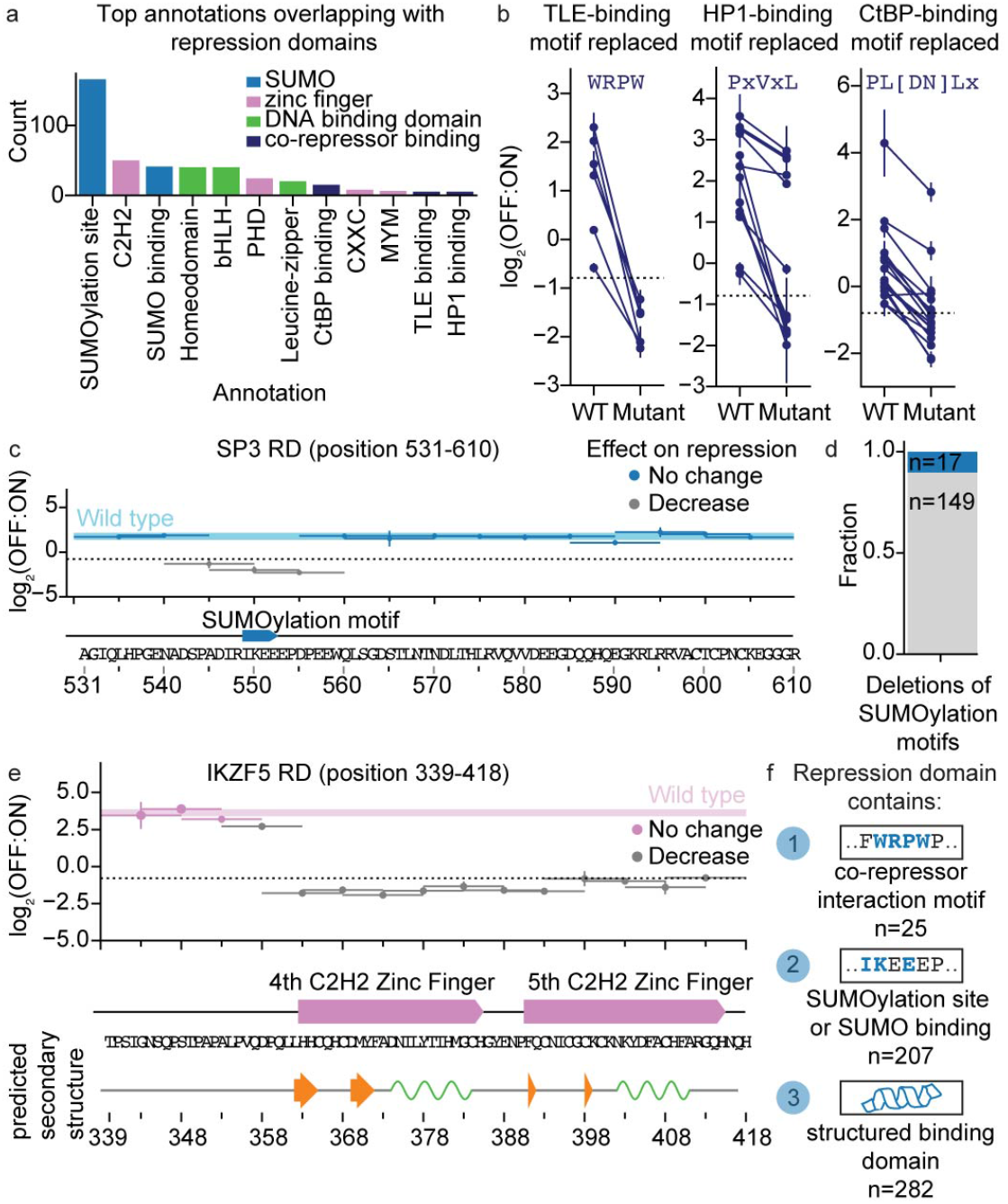
The majority of RD sequences contain either sites for SUMOylation, short interaction motifs for recruiting co-repressors, or are structured binding domains for recruiting other repressive proteins. **a**, Count of repression domains (repressive in both pEF and PGK promoter screens) that overlap annotations from UniProt and ELM (Eukaryotic Linear Motifs). Annotations that had at least 6 counts are shown. **b**, Repression enrichment scores for tiles that contain a co-repressor binding motif (WT) and the co-repressor binding motif replaced with alanines (mutant). TLE-binding: 6 lost all repressive activity upon motif removal. Fraction of non-hit sequences containing motif=0. HP1-binding: 7 lost all repressive activity, 5 decreased, 1 had little change. Fraction of non-hit sequences containing motif=0.002. CtBP-binding: 10 lost all repressive activity, 6 decreased, 1 had little change. Fraction of non-hit sequences containing motif=0.002. 2 biological replicates shown with standard error. **c**, Deletion scan across SP3’s RD. SUMOylation motif is “IKEE” (indicated on the bottom). Blue shaded bar spanning the entire domain length above the threshold represents the WT enrichment score, where the standard error between two biological replicates is represented as the height. Deletions were binned into those that had an effect on repression (gray lines) and those that did not (dark blue lines). **d**, Fraction of deletion sequences containing a SUMOylation motif binned according to their effect on activity (blue=no change on repression relative to WT, gray=decreased repression relative to WT, n=166 total RDs). **e**, Deletion scan across IKZF5’s RD. AlphaFold’s predicted secondary structure (prediction from whole protein sequence) shown below where green regions are alpha helices and orange arrows are beta sheets. **f**, Summary of repression domain functional sequence categories (n indicated in Figure).

Using these filtered tiling data, we annotated repression and activation domains from contiguous hit tiles (**Extended Data Fig. 2d, Methods, Fig. 1b, Supplementary Table 2**). Doing so can accurately identify effector domains previously annotated in UniProt, for example the activation and repression domains in MYB (**Fig. 1b**). Some of the strongest ADs come from gene families with some family members already annotated as activators, such as MYB, ATF, and NCOA, making us more confident our screens returned reliable results. Similarly, some of the strongest RDs come from gene families with some family members already annotated as repressors, such as MBD, KLF, and ZNF gene families (**Fig. 1c-d**). TFs from some gene families, like KLF, ETV, and KMT, contain both strong ADs and RDs, which highlights our results can identify bifunctional transcriptional regulators. In total, 12% of the proteins screened are bifunctional (having both ADs and RDs) and 76% of proteins have at least one effector domain (**Supplementary Table 2**).

In addition, this method allows us to discover previously unannotated effector domains (**Fig. 1e**). For example, we found both a new AD and four new RDs within the DNA demethylating protein, TET2. We validated tens of these new effector domains by individually cloning them, creating stable cell lines, and measuring their effect using flow cytometry after dox-induced recruitment at the minCMV reporter for activation (**Fig. 1f, Supplementary Table 3**) or pEF reporter for repression (**Fig. 1h, Supplementary Table 3**). Doing so, we validated the screen thresholds: all tiles above the thresholds had activity and no tiles below the thresholds had activity (**Fig. 1g**,**i**). In total, 307 of the ADs and 592 of these RDs are new compared to UniProt and previous HT-recruit screen^4^ annotations (**Fig. 1j, Extended Data Fig. 1g**).

Prior screens in yeast have led to the development of a machine learning model (PADDLE^11^) capable of predicting activation levels from sequence alone with an area under the precision-recall curve of 81%. If the sequence properties that drive activation in humans are similar to those in yeast, we would expect PADDLE to predict human ADs with similar accuracy. While PADDLE was able to predict many ADs (70%), the domains that PADDLE predicted to be activating (like the C-terminal AD in CSRNP1) were more negatively charged than the ADs it missed (**Extended Data Fig. 4a**), suggesting that in human cells there are additional non-acidic activator classes compared to yeast.

**Fig. 4.**
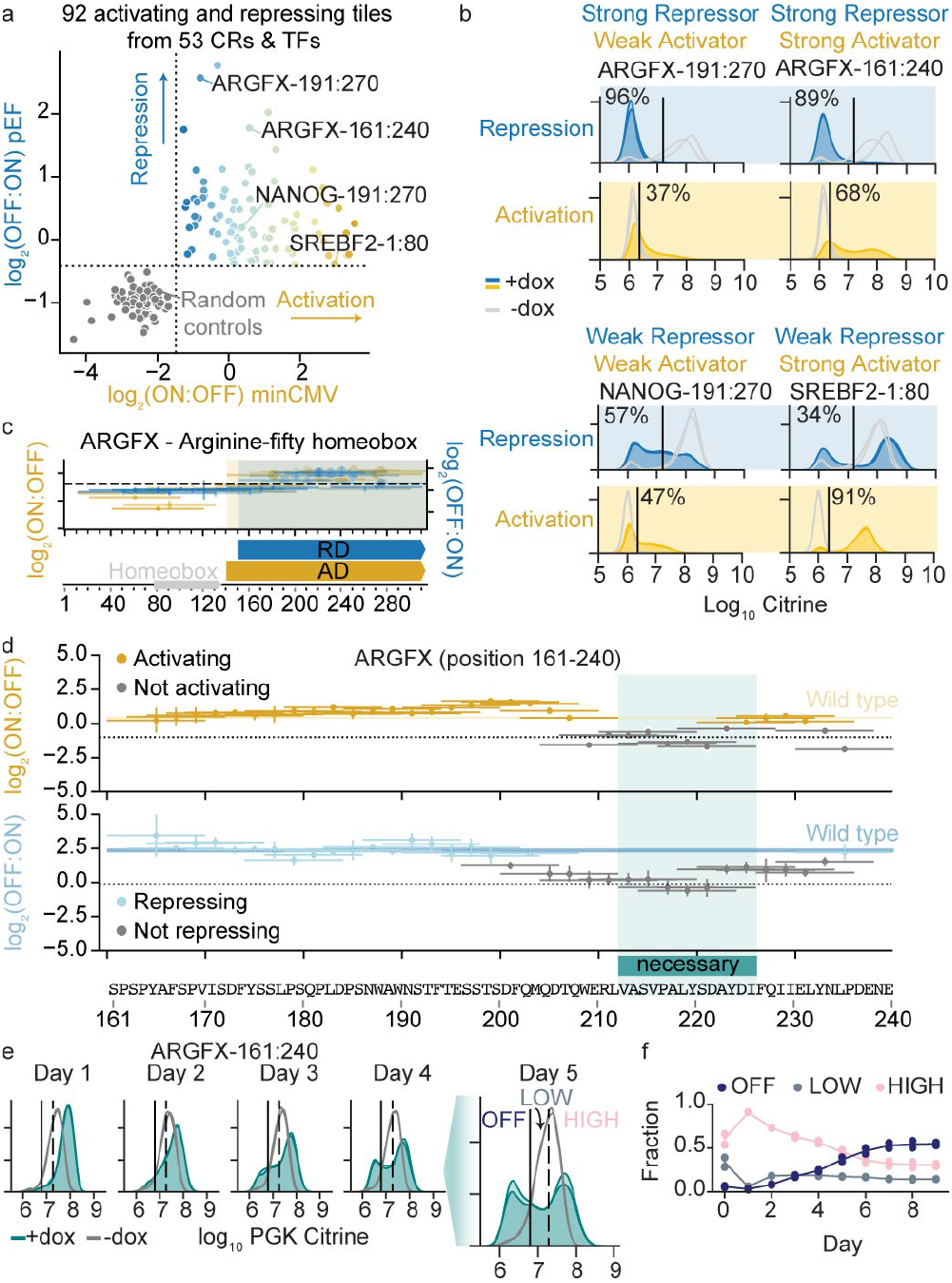
Discovery of bifunctional activating and repressing domains. **a**, Bifunctional tiles were discovered by observing both activation above the hits threshold (vertical dashed line) in the minCMV promoter CRTF validation screen (x-axis) and repression above the hits threshold (horizontal dashed line) in the pEF promoter CRTF validation screen (y-axis). Average across two biological replicates shown for each point. **b**, Individual validations of bifunctional tiles. Untreated cells (gray) and dox-treated cells (colors) shown with two biological replicates in each condition. Vertical line is the citrine gate used to determine the fraction of cells ON for activation and OFF for repression. **c**, Tiling plot for ARGFX. Bifunctional domains are regions where the sequence is both activating at the minCMV promoter and repressing at the pEF promoter. **d**, Deletion scans across ARGFX-161:240 at minCMV promoter (top), and at pEF promoter (bottom). Yellow and blue rectangles represent WT enrichment scores, with the standard error between two biological replicates represented as the heights. The 3 deletions that caused no activation and no repression across both screens are highlighted in teal and the sequence annotated as necessary. **e**, Citrine distributions of bifunctional tile ARGFX-161:240 recruited to the PGK promoter (n=2). Left vertical gate was used for measuring the fraction of cells OFF to its left. Right vertical gate was used for measuring the fraction of cells HIGH to its right. The fraction of LOW cells were measured as the cells in between both gates. **f**, Fraction of ARGFX-161:240 cells OFF (navy), LOW (gray), and HIGH (pink) over time (2 biological replicates plotted with the average plotted as a line).

Observed repression at the pEF promoter could reveal sequences that universally repress gene expression; alternatively, this behavior may depend on the promoter to which tiles are recruited. To distinguish between these scenarios and because there are no other comprehensive studies to reference our results to, we decided to determine how many of these tiles are repressor hits at a different constitutive promoter. We performed a new screen of the CRTF tiling library at the PGK promoter. While this promoter is weaker, we were able to separate the silent and active cells by magnetic separation (**Extended Data Fig. 4b**) and observed good reproducibility across two replicates (**Extended Data Fig. 4c**). 92% of the hit tiles that showed up in the pEF and PGK screens also showed up as hits in the pEF validation screen (**Extended Data Fig. 4d**), suggesting higher confidence results when combining both screens. We called RDs from contiguous hit tiles in the PGK data (**Extended Data Fig. 4e**). Across the two repressive screens (at pEF and PGK), we found a total of 3,900 repressor domains, noting that some of these domain boundaries are overlapping. Taking the maximum tile’s enrichment scores within each RD revealed 715 RDs were shared across both screens (**Extended Data Fig. 4f**). Together, these results suggest that at the 80 aa scale there are more sequences across the CRs and TFs that can work as repressors versus activators.

### Activation domain sequence characteristics

ADs have been classified by the abundance of particular amino acids such as acidic (D, E), glutamine-rich (Q), and proline-rich (P) sequences^10,27^. Acidic residues have been shown to be essential for function in all yeast activation domains^11^ and some human ADs^15^. Certain human ADs have compositional biases that are not present in other organisms, often containing stretches of single amino acid homotypic repeats^28^ (i.e. QQQQ). Additionally, some human ADs are enriched in particular hydrophobic residues - aromatics (W, F, Y) and leucines (L), that are important for function in that context^15^. It remains unclear how many human ADs fall under each of these categories - acidic, compositionally biased, or hydrophobic, if these categories are indeed distinct from one another, and if these amino acids that are enriched have functional significance for the ADs in those categories. Specifically, would activation be lost if we deleted or mutated these amino acids?

Our new large set of activating tiles provides a great opportunity to systematically quantify the prevalence of each of these sequence properties within human ADs. Every activating tile contained at least one aromatic or leucine residue, and nearly every tile contained at least one acidic residue (**Fig. 2a**). Moreover, 45% of activating tiles contained a compositional bias (**Fig. 2a**), where serine (sequences from NR4A and NFE2 families) and proline (sequences from FOX family and P53) were the most abundant. Given that several ADs have been categorized by their glutamine content, we were surprised to find very few glutamine-rich ADs across all the CRs and TFs (sequences from SMARCA family and TRERF1, **Supplementary Table 4**). Consistent with these observations, when we normalize the amino acid frequencies in the AD sequences by the amino acid counts in the non-hit sequences, we observe an enrichment in certain hydrophobic, acidic, serine, and proline residues (**Fig. 2b**).

To determine which amino acid types among these enrichments are necessary for activation and to find the necessary sequences within each activating tile, we took a deletion scanning approach, which others have used to identify necessary sequences in yeast ADs^29^. First, by performing scanning deletions (15 aa each) across 24 UniProt annotated ADs that had activity at the minCMV promoter in K562, we found that while most (61%) deletions do not affect activation, in the majority of these ADs (20/24) we found at least one deletion that was well-expressed and could abolish activator function (**Supplementary Table 1**). In order to validate that this deletion scanning approach returned residues that were necessary for activity, we compared our deletion scan data from P53 to UniProt annotations and found the minimized necessary sequences to be residues 20-22 (DLW) within one region and residue W52 within another region, corresponding to UniProt-annotated TAD I and TAD II, respectively (**Extended Data Fig. 5a**). Furthermore, individual validations confirmed the complete loss of activity when deletions including these residues were tested (**Extended Data Fig. 5b**).

Confident in our deletion scan approach, we designed a second library of 10 aa deletions across the maximum activating tile from each AD, resulting in 304 total deletion scans (**Supplementary Table 4**). We measured activation scores for all 12,320 members of this library using the minCMV reporter and HT-recruit workflow described in **Fig. 1a** (**Extended Data Fig. 5c-d**). We FLAG-stained for protein expression (**Extended Data Fig. 5e-f**) and filtered out mutants that were poorly expressed. Across each of these expression-filtered deletion scans we first binned deletions into those that had an effect on activation and those that did not (**Fig. 2c**). Using these data, we can identify which of the amino acids that contributed to the compositional bias are important for function: for example, while NFAT5’s AD has a patch of 4 serines near the C-terminus, deleting those residues had no effect on activation (**Fig. 2c**). We highlight similar examples for stretches of prolines and glutamines that are not essential for activation (**Extended Data Fig. 6a**). Applying this analysis to all ADs containing a homotypic repeat, and after removing all poorly-expressed deletions, we find homotypic repeats of certain hydrophobic residues like glycine, alanine and leucine were equally found in deletions that had no effect on activation and in deletions that decreased activation (**Fig. 2d**). However, serine, proline, acidic and glutamine homotypic repeats were more often found in deletions that had no effect on activation than in deletions that decreased activation (**Fig. 2d**). Therefore, homotypic repeats of these amino acids are generally not necessary for activation.

The deletion scans also allow us to identify the necessary sequence for activation of each tile: sequences that, once removed, completely abolished activation (**Fig. 2c**). We were able to annotate at least one necessary sequence (median length=10 aa) in the majority (69%) of our screened ADs, and most (61%) ADs have multiple necessary sequences, supporting the idea that ADs are composed of multiple small linear binding motifs (**Fig. 2c, Supplementary Table 4**). Nearly every necessary sequence (96%) contained a W, F, Y or L.

In order to validate this enrichment of specific hydrophobic residues, we rationally designed mutant libraries where we systematically replaced every amino acid of a particular type within the sequence with alanines (**Supplementary Table 4**). Replacement of all W, F, Y or Ls with alanine (range: 3-24 aa replaced/80 aa tile, median=10 aa) in all our activating tiles resulted in a total loss of activation (**Fig. 2e**). The one exception that remained active was within DUX4, and the mutation did in fact make it weaker (**Extended Data Fig. 6b**). This systematic loss of activation was not due to a decrease in protein expression, as measured by FLAG staining (**Extended Data Fig. 6c**). This means all 732 tested tiles from 258 proteins with ADs require some aromatic or leucine residues to activate.

We next wanted to follow up more on the acidic sequences, so we replaced all acidic residues with alanine in the entire set of activating tiles (not just the few that had a compositional acidic bias). Surprisingly, more than half of the acidic mutants had reduced expression (**Extended Data Fig. 6c**). These results suggest acidic residues increase protein levels, at least in the context of transcriptional activators. Of the remaining 247 well-expressed activating tile mutants, the majority of mutants lost the ability to activate (**Fig. 2f**, n=196). 33 mutants decreased their activities upon mutation, and only 18 mutants had no change in activation, where some in fact increased (**Supplementary Table 4**). The activator tiles that depended on acidic residues came from a wide range of TF families, including E2Fs and GRHLs and the classical example acidic activator ATF4 (GCN4’s mammalian homolog). Some of the sequences that do not require acidic residues came from SMARCAs, TET2, PLAG1s, and every paralog from the EYA family that had an AD. These mutants with no change in activity had significantly fewer acidic residues than the tiles whose mutants had a decreasing effect (**Extended Data Fig. 6d**), supporting the idea that acidic ADs are not the only class of human ADs.

Intrigued by what other compositional biases could be functional in human ADs, we next tested the necessity of other frequently-appearing amino acids. We replaced compositionally biased amino acids with alanine. For the few activation tiles that contained glycine-rich and glutamine-rich sequences, there were fewer than 5 mutants that expressed well as measured by FLAG (**Supplementary Table 4**), so we excluded these from further statistical analyses. Consistent with the results above, all tiles with leucine compositional biases lost activity once mutated, and the few tiles with acidic biases lost activity once mutated (**Fig. 2g**). Removal of serine and proline compositional biases had more mild effects: the vast majority of mutants still had activity (**Fig. 2g**, top), even though the strength of activation decreased for a subset of them (**Fig. 2g**, bottom).

Wanting to follow up more on the compositionally biased tiles that decreased activity upon compositional bias removal (**Fig. 2g**), we next wondered if it was the homotypic repeats themselves that explained this loss in activity or if a subset of compositionally biased residues overlapped important co-activator binding motifs. To answer whether the placement of serines, prolines, and acidic residues within the sequence were more important than their overall abundance, we analyzed the set of deletion necessary sequences from the compositionally biased activating tiles that lost activity upon bias removal (**Fig. 2g**, bottom). For each compositional bias type, the majority of necessary sequences also contain a W, F, Y, or L (**Fig. 2h**).

In summary, sequences that are necessary for activation consist of certain hydrophobic amino acids (W, F, Y, and/or L) that are interspersed with either acidic, proline, serine, and/or glutamine residues (**Fig. 2i, Extended Data Fig. 6e**).

### Repression domain sequence characteristics

Repressing tile sequences have significantly more secondary structure than activating tile’s (**Extended Data Fig. 7a**). Therefore, we needed to take a different approach for understanding the sequence characteristics of RDs. Instead of looking at RD sequence compositions, we first set out to classify the RDs by their potential mechanism. We used the ELM database to search for co-repressor interaction motifs (**Methods**), and UniProt to search for domain annotations. We observe 72% of the RDs overlap diverse annotations, such as sites for SUMOylation, zinc fingers (C2H2, PHD, CXXC, MYM), SUMO-interacting motifs, co-repressor binding motifs (CtBP-, HP1-, TLE-binding), DNA binding domains (Homeodomain DBDs, consistent with previous results^4^), and dimerization domains (bHLH, Leucine-zipper) (**Fig. 3a**). To address whether these annotated sequences are necessary for repression, we rationally designed mutant libraries that systematically replaced sections of 1,313 RDs (**Supplementary Table 5, Methods**) and screened this RD mutant library using the pEF reporter and workflow described in **Fig. 1a** (**Extended Data Fig. 7b-c**). We stained for protein expression (**Extended Data Fig. 7d-e**) and filtered out mutants that had low FLAG enrichment scores.

First, we systematically searched and replaced the co-repressor interaction motifs with alanine to test their contribution to activity (**Fig. 3b**). The TLE-binding motif, WRPW, appears exclusively in the C-terminal RDs of the HES family and all tiles containing this motif were repressive (**Extended Data Fig. 7f**). All tested motifs were necessary for repression (**Fig. 3b**, left). The HP1-binding motif, PxVxL, was necessary or contributed to repression in the majority of the tiles containing it (12/13 tiles with decreasing effects **Fig. 3b**, middle). CtBP’s binding motif, Px[DENS][LM]x, and the SUMO interaction motif, -fxff-(non-covalent binding site to SUMOylated proteins, found in co-repressors that promote heterochromatin formation such as SETD1a), are both relatively more flexible than the former two motifs and therefore appeared in more RDs. However, in many RDs, they are not essential for function, as their deletion does not decrease repression (**Extended Data Fig. 8a-b**). We found that a more refined CtBP motif of PL[DN]Lx explained the majority of tiles that lost activity upon mutation (16/17 tiles **Fig. 3b**, right). Altogether, 94% of the 36 repressing tiles with a co-repressor associated motif (TLE-, HP1-, or CtBP-binding) decreased in repression strength when the motif was mutated, while 72% of 113 SUMO interaction motif-containing repressing tiles were similarly sensitive to mutation (**Extended Data Fig. 8b**).

We were intrigued by the many RDs that contain a SUMOylation site (site for covalent conjugation of a SUMO domain) (**Fig. 3a**). The ELM database classifies SUMOylation sites with the search pattern φKxE. Because this motif is short and relatively flexible, some non-hit sequences (12.3%) also contain SUMOylation motifs. In order to investigate whether SUMOylation sites within non-hit sequences are functional, we first used the AD deletion scan data. Deleting a SUMOylation motif within ADs rarely decreased activation (**Extended Data Fig 8c**). Next, we asked if these motifs are functional in RDs using the same deletion scanning approach (**Supplementary Table 5, Fig. 3c**). For example, residue K550 in the SP3 protein is a SUMOylation site and has been shown before to be important for repression^30^; indeed we also find the SUMOylation site to overlap with the region essential for repression for this RD of SP3 (**Figure 3c**). In a similar manner, we find SUMOylation motifs are important for the repression of at least 149 out of the 166 RDs where they are found (**Fig. 3d, Supplementary Table 5**). This result is concordant with our previous finding that a short 10 amino acid tile from the TF MGA, which contains this SUMOylation motif, IKEE, is itself sufficient to be a repressor^4^. While the role of this modification in repression still needs to be better understood, SUMOylation of certain TFs, such as FOXP1 (which also shows up as a necessary region in our measurements, **Supplementary Table 5**), has been shown to promote repression via CtBP recruitment^31,32^. Our results suggest the pervasive role, across over a hundred TFs, that SUMOylation plays in repression.

We next used our deletion scan data to gain better resolution of the region within RDs overlapping dimerization domains, such as basic helix-loop-helix domains (bHLHs). Within bHLHs, the basic region binds DNA, and mutations in the HLH region are known to impact dimerization^33^. Deletion scans across tiles that overlap HLH domains reveal part of helix 1, the loop region, and helix 2 are necessary for repression (**Extended Data Fig. 8d**). The majority of RDs that overlap HLHs can be classified as Class II, tissue specific dimerization domains that can either be activating or repressing depending on the context^33^ (**Extended Data Fig. 8e**). Our data suggests many Class II bHLHs can function as RDs. This does not exclude the possibility bHLHs can also function as ADs, but we only observe NEUROG3’s bHLH activate the minCMV promoter, suggesting there is promoter specificity to activation of HLH domains.

Many RDs overlap annotated zinc fingers (n=124), and some specifically overlap C2H2 zinc fingers (n=50, compared to only 3 ADs that overlap C2H2 zinc fingers) (**Fig. 3a**). We wondered if the C2H2 domain itself or the protein sequence flanking it was responsible for repression. For example, REST’s zinc finger directly recruits the co-repressor coREST^34^, and indeed REST deletions that had no effect on repression (pink) corresponded to the disordered region just outside of the zinc finger, and deletions necessary for repression (gray) corresponded to the zinc finger structural fold (**Extended Data Fig. 8f**).

In addition to binding DNA and directly binding co-repressors, zinc fingers dimerize with other zinc fingers^35^. We reasoned some zinc fingers could cause repression by binding to other zinc finger domains within endogenous repressive proteins. Support for this indirect recruitment of repressive TFs via zinc fingers comes from the IKZF family where the N-terminus of some members, such as IKZF1, directly recruits CtBP^36^, while the C-terminal zinc fingers bind other IKZF family members^37^. Indeed, we recover the N-terminal repressive domains in IKZF1, and the associated sequence contains a CtBP binding motif (**Extended Data Fig 8g**). In addition, all IKZF family members show C-terminal RDs that overlap the last two zinc fingers (**Extended Data Fig 8g**). These two zinc fingers are both necessary for repression in IKZF5 (**Fig. 3e**) and in all tested family members (**Extended Data Fig 8h**), and therefore likely dimerizes with the IKZFs that recruit CtBP. While in general zinc fingers are well-known DNA binding domains, our data expands the list of zinc finger sequences that are likely protein binding domains to other repressive TFs (**Supplementary Table 5**).

In summary, repression domains can be categorized by their sequence properties in the following way: (1) domains that contain short, linear motifs that directly recruit co-repressors, (2) domains that contain SUMO interaction motifs or can be SUMOylated and most likely recruit co-repressors through the conjugated SUMO domain, or (3) structured repressive protein binding domains that can recruit co-repressors or other repressive TFs (**Fig. 3f, Extended Data Fig 8i**).

### Bifunctional activating and repressing domains

Transcriptional proteins are often categorized as activating, repressing, or bifunctional^8^. Bifunctionality is when the protein activates some promoters but represses others^38^. There are 248 bifunctional CRs & TFs that have both an AD and RD (such as in **Fig. 1b, Supplementary Table 2**). Additionally, we observe bifunctionality at the domain level, wherein the same 80 aa tile both activated a minimal promoter and repressed a constitutive promoter (**Fig. 4a-c, Supplementary Table 6**). We wondered if the 92 bifunctional domains we discovered appear in specific TF families and found many are within homeodomain TFs (**Extended Data Fig. 9a**).

We individually validated bifunctional domains by flow cytometry, and confirmed doxycycline-dependent activation of the minCMV and repression of the pEF reporter genes for all tested domains (**Fig. 4b, Supplementary Table 3**). Some domains have both weak repression and activation, like the tile from NANOG (**Fig. 4b, Extended Data Fig. 9b**). Some domains are stronger activators than repressors, some stronger repressors than activators, and other domains show both strong activation and repression (**Fig. 4b**). Together, the screen and validations demonstrate the CRTF tiling library can be screened at multiple promoters to uncover bifunctional domains.

We hypothesized most bifunctional domains are similar to bifunctional proteins (as in **Fig. 1b**), composed of smaller activating and repressing regions at independent locations. To address whether the same exact sequence could be responsible for activation and repression we did a deletion scan across all 92 bifunctional domains at the minCMV and pEF reporters (**Supplementary Table 6, Extended Data Fig 9c-f**). These deletion scans revealed that some bifunctional tiles, including ones in NANOG (**Extended Data Fig 9g**), have independent activating and repressing regions. In contrast, in other tiles, the same amino acids are necessary for both activation and repression: for example, a single 14 aa region mediated both the strong activation and repression for ARGFX tile 16 (**Fig. 4d**). Similarly, the same 14 aa region is necessary for both activities for LEUTX, a TF in the same gene family as ARGFX (**Extended Data Fig 10a-b**). In summary, a region as small as 14 amino acids can be necessary for both activating and repressing activities, and as many as 69 other bifunctional domains similarly contain single regions that are necessary for both activities (**Extended Data Fig 10c**).

In individual validations that measured activation over a time course, bifunctional ARGFX tile 16 (**Figure 4b**) was stronger at activating the minCMV promoter at the day 1 time point compared to day 2 (when the screen was measured) (**Extended Data Fig. 10d**), and in fact, the activated promoter slowly silenced upon further recruitment to day 4. Intrigued by these dynamics, we tested several bifunctional tiles at a promoter that has intermediate levels between minCMV and pEF, to see which direction they would tune transcription. Surprisingly, when we recruited ARGFX tile 16 to the intermediate promoter, we observed both a highly activated population of cells and a repressed population of cells after 5 days of recruitment (**Fig. 4e**). Similar to the minCMV promoter, most cells initially increased in expression at day 1, then a subpopulation of cells silenced while another remained high (**Fig. 4e-f**). Other bifunctional tiles recruited to the PGK promoter, from FOXO1 and ARGFX, led to similar dynamics that start with activation and eventually end in a split of the cell population into silenced cells and cells that continue to express (**Extended Data Fig. 10e**). These tiles, like ARGFX tile 16, had overlapping regions that are necessary for both activities (**Supplementary Table 6**). However, not every bifunctional tile that activates minCMV and represses pEF has bifunctional activity at the PGK promoter (**Extended Data Fig. 10e**): for example, NANOG and KLF7 tiles do not significantly change expression of the PGK promoter. These tiles, in contrast, have independent activating and repressing regions (**Extended Data Fig. 9g, Supplementary Table 6**). In summary, some bifunctional tiles that independently activate and repress different promoters are bifunctional even at a single promoter and can dynamically split a cell population into high- and low-expressing cells.

## Discussion

A systematic understanding of how transcriptional proteins function in human cells is needed to make medical advances. When a new transcriptional protein is sequenced, homology models robustly identify the DNA binding domain locations, but are unable to predict where the effector domains are^39^. Compared to DNA binding domains, many effector domain sequences are poorly conserved and do not align well with one another in a multiple sequence alignment. As a result, we do not have nearly as robust nor as comprehensive of predictors or sequence patterns for finding effector domains within protein sequences, and thus need high-throughput experimental approaches for discovering them.

Here, we report the most comprehensive measurements to date of human transcriptional effector domains. Via high-throughput tiling screens combined with systematic deletion scans and rational mutagenesis, we collectively assigned transcriptional effector domains to 76% of the CRs and TFs screened and comprehensively dissected the sequence properties that are necessary for activation and repression.

The sequences that are necessary for function in ADs consist of certain hydrophobic amino acids (W, F, Y, or L) that are interspersed with either acidic, proline, serine or glutamine residues (**Fig. 2i**). Although prior work has shown homopolymeric stretches of glutamine and proline are sufficient to activate a weak synthetic reporter^28^, we find only OCT4’s AD has proline repeats that are necessary for activation. In fact, the majority of glutamine and proline repeats within ADs of the human CRs and TFs are not part of the sequence necessary for activation. While these homotypic repeats might still be important for other effects within the full-length TFs, such as solubility or nuclear localization, our data suggests they are generally dispensable parts of AD sequences. In addition to the acidic exposure model, our data suggests additional ways human ADs promote hydrophobic exposure, where serines could functionally mimic acidic residues when phosphorylated, and prolines could promote exposure by their intrinsic disorder. Furthermore, ADs contain certain hydrophobic amino acids, but our data suggest those residues can be arranged in many ways, interspersed with serine, proline, and/or acidic residues. Unlike RDs, we did not find any AD motifs, other than the previously reported LxxLL which appeared in 41 activating tiles (**Supplementary Table 4**). AD grammar flexibility might be related to their promiscuity in binding, where many ADs have been shown to bind to more than one co-activator target^40^, likely because co-activators are a scarce resource in the cell^41^. ADs lacking motifs, or having little grammar, might impart their flexibility by binding several different co-activators. In order to improve our understanding of ADs, it will be important to take the next step and dissect how their sequence composition relates to their binding to co-activators.

Strong sequence enrichment patterns across families have never been observed for RDs^8^. This observation is likely due to the fact that there are many distinct functional categories of RDs. Indeed, 514 of our RDs overlap diverse functions, including co-repressor binding motifs, SUMO interaction motifs, and structured binding domains. It has been shown before that the presence of the SUMO-1 domain alone is sufficient to cause repression^30^, and some well-characterized RDs contain SUMOylation sites^32^. Here, we find hundreds of RDs that contain SUMOylation sites and show that repression activity is lost upon deletion of these sites for the majority (>90%) of these RDs (**Figure 3d**). One mechanism by which SUMOylation leads to repression is by recruitment of SUMO interaction motif-containing co-repressors (**Extended Data Fig. 8i**). Supporting this, mutagenesis of SUMO interaction motifs within our set of RDs, for example in SETD1a, led to a reduction of repression in 81/113 tested tiles (**Extended Data Fig. 8b**). An alternative hypothesis for SUMOylation-mediated repression is that SUMOylation affects the TF’s localization within the nucleus towards regions associated with heterochromatin^30,32^, but more investigation into the mechanism of each SUMOylated RD will be needed.

Zinc finger domains were originally identified as DNA binding domains, yet many of these domains are also protein binding domains^35^. Very few examples of transcriptionally repressive zinc finger domains existed before this study. Here, our data suggest zinc finger domains are a prevalent repression sequence, which are necessary for the repression of hundreds of domains. While a handful of zinc fingers have been shown to interact with corepressors or repressive partner TFs, for most of them, their partners remain to be found.

By systematically measuring both activation and repression of the same library, we were able to find effector domains that can perform both roles. While bifunctional TFs that contain separate activator and repressor domains have previously been observed^42–44^, to our knowledge, this is the first observation of bifunctional domains that are capable of simultaneously enhancing and silencing expression from a single promoter. Deletion scan data revealed activating and repressing regions within these bifunctional domains can be very close to one another (less than 80 amino acids apart), and even overlapping in the majority of domains. Previous observations of master transcriptional regulators activating some genes and repressing others, such as NANOG^45,46^, might be explained by this protein’s bifunctional domain. We find it interesting that bifunctional domains are most commonly found in the homeodomain family of TFs (**Extended Data Fig. 9a**). Many homeodomain DBDs are not sufficiently specific to bind DNA on their own and thus compensate by either having multiple motifs or multiple proteins helping the homeodomain bind its enhancer^47,48^. Therefore, the direction of the bifunctionality (whether the gene gets activated or whether it becomes repressed) might be tuned by the DBD’s motif within an enhancer/silencer. Evidence for this hypothesis has been shown for the bifunctionality of the homeodomain TF CRX where observing repression, in addition to activation, depends on the number of CRX binding motifs in a synthetic context or the presence of other TF binding motifs in a genomic context^49,50^. The functional difference between a bifunctional protein’s silencer or enhancer sequence might simply be explained by the bifunctionality of the effector domain and the ratio of recruited activating and repressing complexes, where more binding motifs will lead to repression. It will be interesting to determine how the CRs and TFs with bifunctional domains that lead to a pulse of activation followed by repression of the gene affect development and patterning.

Although we have acquired quite an extensive dataset, there is still more to be discovered by using the same approach and libraries, and performing these high-throughput measurements in other cell types and under different signaling conditions. Nevertheless, this is one of the largest high-throughput assays for protein function performed in human cells to date, where we followed up with smaller high-throughput validations and protein expression measurements in order to produce a high quality and comprehensive dataset, moving one step closer to proteome-wide functional screening of protein domains.

Now, this catalog can be used for improving sequence prediction models of transcriptional effector domains, understanding molecular principles and the possible effects of CR and TF disease mutants, and engineering better synthetic transcription factors and CRISPR systems^51^. We anticipate this resource will enable exploration of uncharted functional genomic studies.

## Extended Data Figures

**Extended Data Fig. 1.**
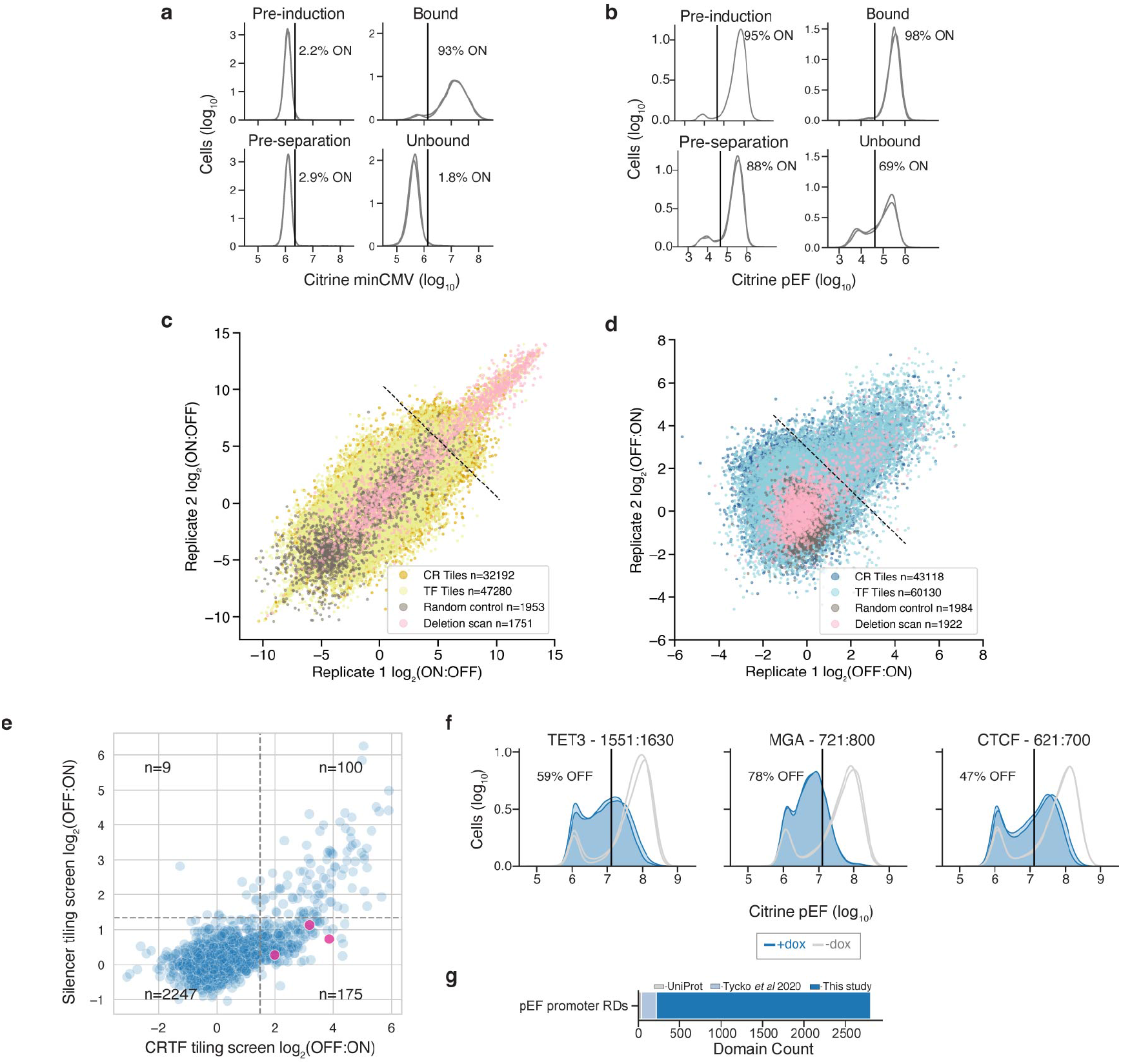
CRTF tiling screen’s separation purity, reproducibility, and validation. **a**, Flow cytometry data showing citrine reporter distributions for the minCMV promoter screen on the day we induced localization with dox (Pre-induction), 2 days later on the day of magnetic separation (Pre-separation), and after separation using ProG DynaBeads that bind to the surface synthetic marker (Bound). Overlapping histograms are shown for 2 separately transduced biological replicates. The average percentage of cells ON is shown to the right of the vertical line showing the citrine level gate. A total of 1,000 ng/mL dox was added each day of dox treatment. **b**, Citrine reporter distributions for the pEF promoter screen (n=2). Pre-separation was after 5 days of dox treatment. **c-d**, Biological replicate screen reproducibility (pearson r^2^=0.78 for minCMV and r^2^=0.19 for pEF hits). **e**, Comparison between repression enrichment scores of tiles that were screened in the CRTF tiling pEF screen (x-axis) and previous Silencer tiling screen (y-axis)^4^. Dashed lines are the hits thresholds for each screen. Tiles were identical with a 1 amino acid register shift (as Silencer library tiles included an initial methionine absent from the CRTF tiling library). Pink dots are tiles that were individually validated in f. **f**, Citrine reporter distributions of individually validated CRTF tiling pEF screen hits that were not identified within the Silencer tiling screen. **g**, Counts of RDs annotated from tiles that were hits in the pEF promoter screen. Domain counts that are new (dark blue), overlap UniProt annotations (gray), or overlap prior HT-recruit screen results^4^ (light blue). Total of 2,803 domains, where 2,585 are new when recruited at the pEF promoter.

**Extended Data Fig. 2.**
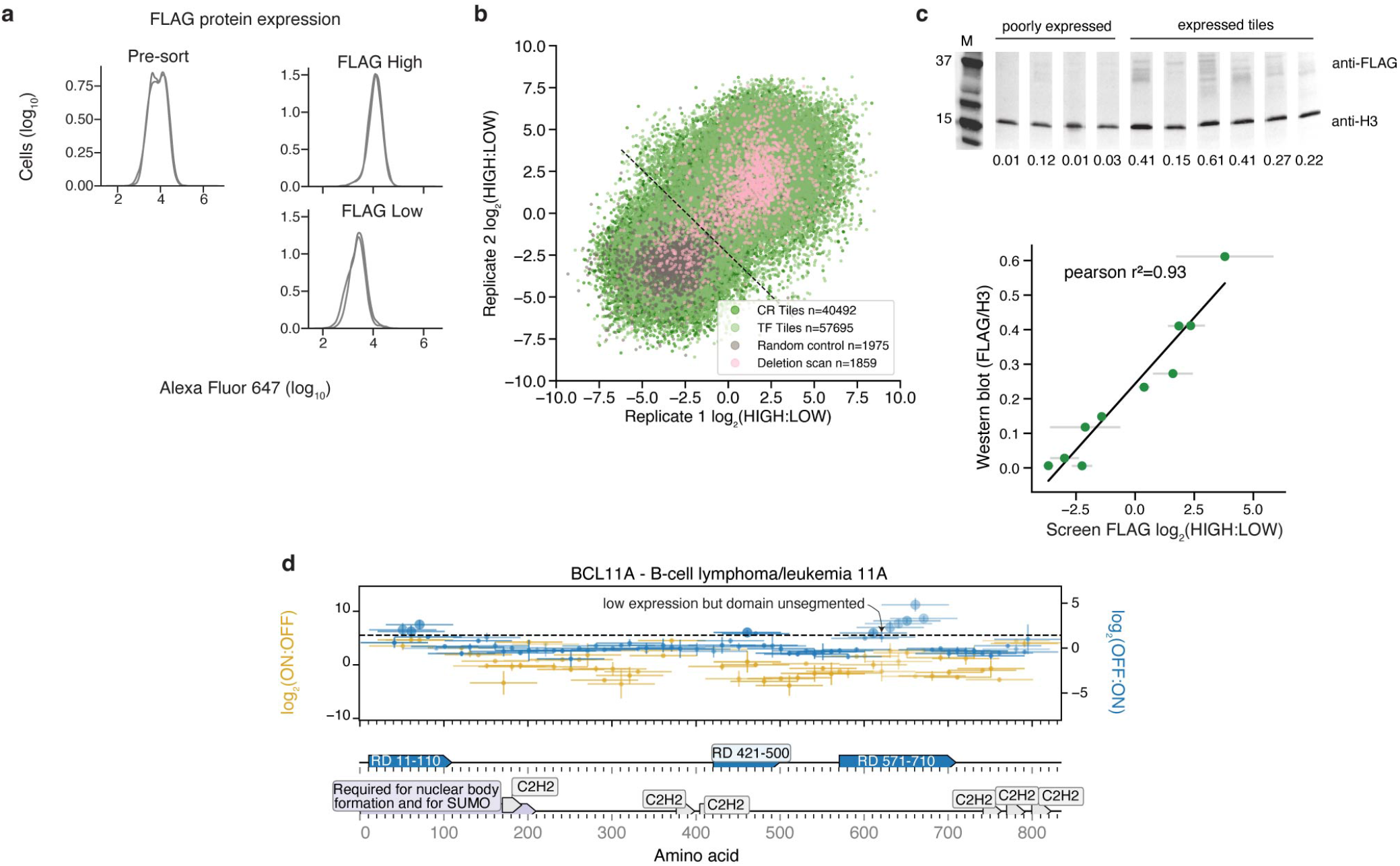
CRTF tiling FLAG protein expression screen separation purity, reproducibility, validation, and example of how the data were used. **a**, Alexa Fluor 647 distributions from anti-FLAG staining of the CRTF tiling library in minCMV promoter reporter cells (n=2). **b**, Biological replicate screen reproducibility. **c**, Validation of expression level for a panel of tiles. Expression level was measured by western blot with an anti-FLAG antibody. Anti-histone H3 was used as a loading control for normalization. Levels were quantified from all bands in each lane using ImageJ. Superfluous lanes from the gel are cropped out and the relevant lanes are shown consecutively with white lines between each lane. Comparison between high-throughput measurements of expression and western blot protein levels (r^2^=0.93). **d**, Tiling plot for BCL11A. Example of a domain that was annotated at position 571-710. This domain had a low expression tile in the middle but the domain was left unsegmented. See more about how domanis were called in Methods.

**Extended Data Fig. 3.**
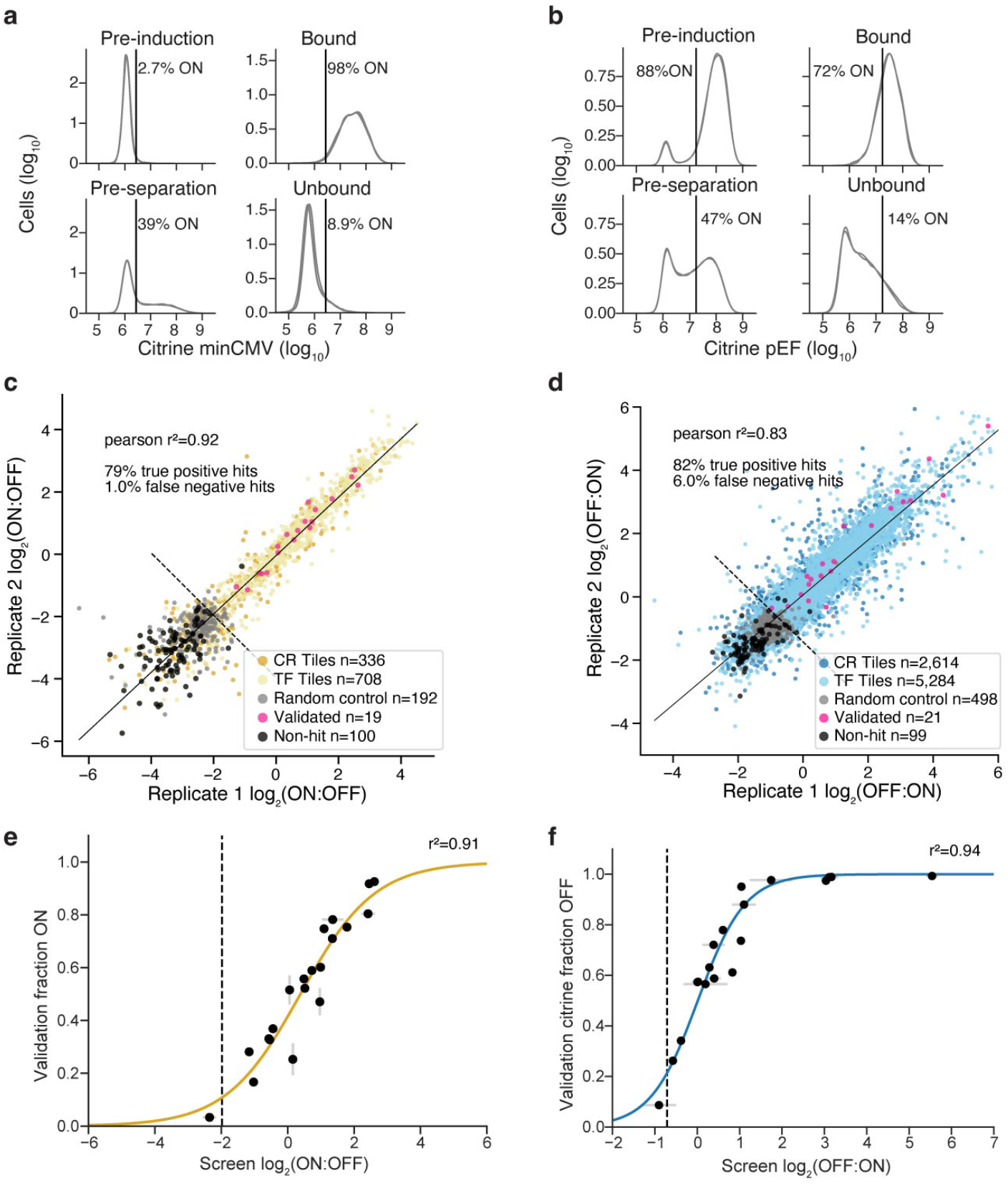
CRTF tile hits validation screen’s separation purity, reproducibility, and validation. **a**, Flow cytometry data showing citrine reporter distributions for the minCMV promoter screen on the day we induced localization with dox (Pre-induction), 2 days later on the day of magnetic separation (Pre-separation), and after separation using ProG DynaBeads that bind to the surface synthetic marker (Bound). Overlapping histograms are shown for 2 biological replicates. The average percentage of cells ON is shown to the right of the vertical line showing the citrine level gate. A total of 1,000 ng/mL dox was added each day of dox treatment. **b**, Citrine reporter distributions for the pEF promoter validation screen (n=2). Pre-separation was after 5 days of dox treatment. **c-d**, Biological replicate screen reproducibility. **e**, Comparison between individually recruited measurements and minCMV promoter validation screen measurements with logistic model fit plotted as solid line (r^2^=0.91, N=20). **f**, Comparison between individually recruited measurements and pEF promoter validation screen measurements with logistic model fit plotted as solid line (r^2^=0.94, N=19).

**Extended Data Fig. 4.**
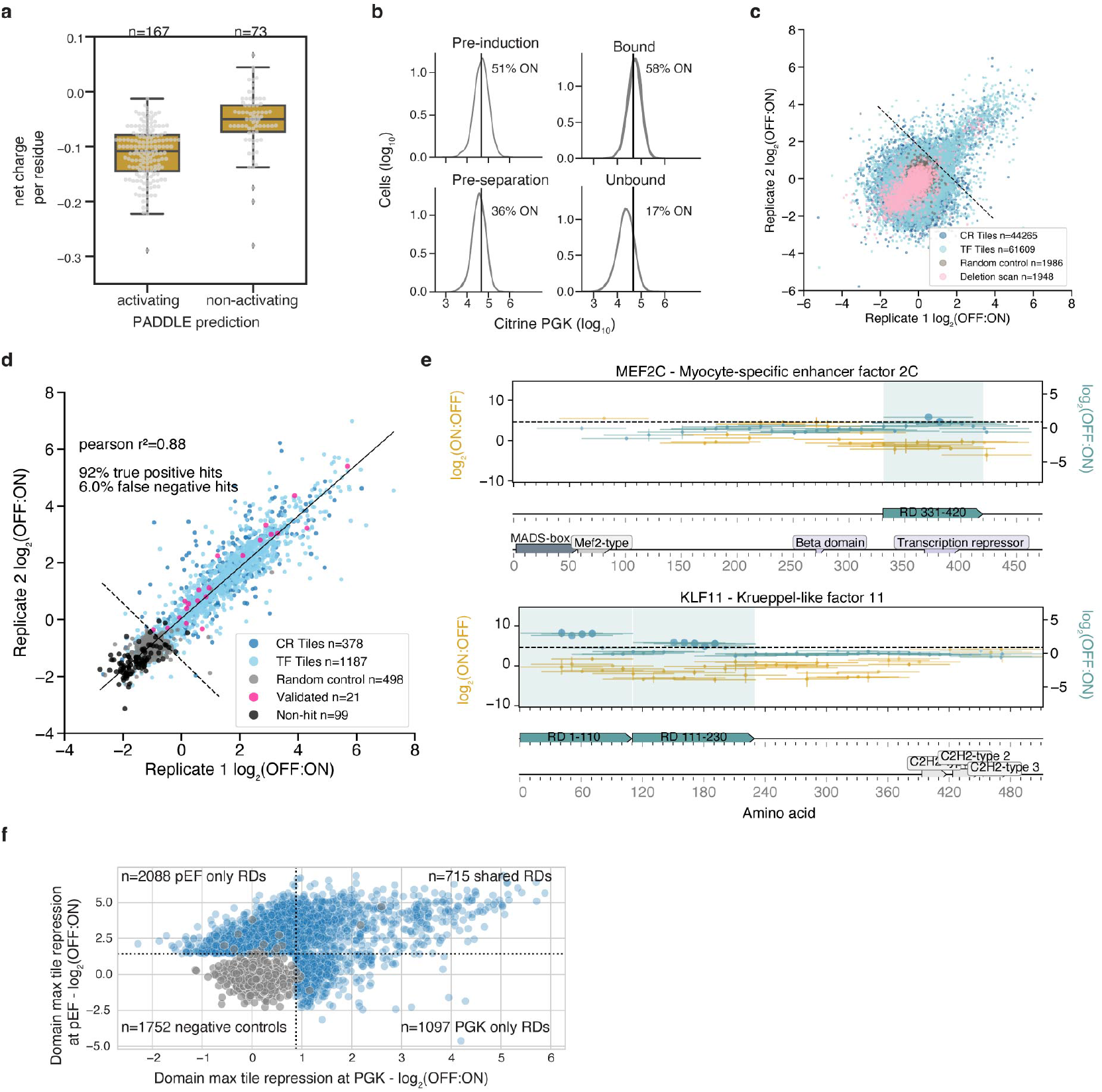
Validations of CR & TF effector domains. **a**, Net charge per residue distributions (calculated by CIDER^52^) of activation domains identified by HT-recruit compared to their PADDLE-predicted function^11^ (Mann-Whitney p-value=1.39e-15). **b**, Flow cytometry data showing citrine reporter distributions for the PGK promoter screen on the day we induced localization with dox (Pre-induction), 5 days later on the day of magnetic separation (Pre-separation), and after separation using ProG DynaBeads that bind to the surface synthetic marker (Bound). Overlapping histograms are shown for 2 biological replicates. The average percentage of cells ON is shown to the right of the vertical line showing the citrine level gate. A total of 1,000 ng/mL dox was added each day of dox treatment. **c**, Biological replicate PGK promoter screen reproducibility (pearson r^2^=0.27 for hits). **d**, Validation screen biological replicate reproducibility of tiles that were hits in both the PGK and pEF promoter screens. **e**, Tiling plots for MEF2C and KLF11. PGK repression domains annotated in teal. **f**, Comparison of each repression domain’s max tile repression scores in PGK (x-axis) and pEF promoter screen (y-axis). Dashed lines are the hits thresholds for each screen.

**Extended Data Fig. 5.**
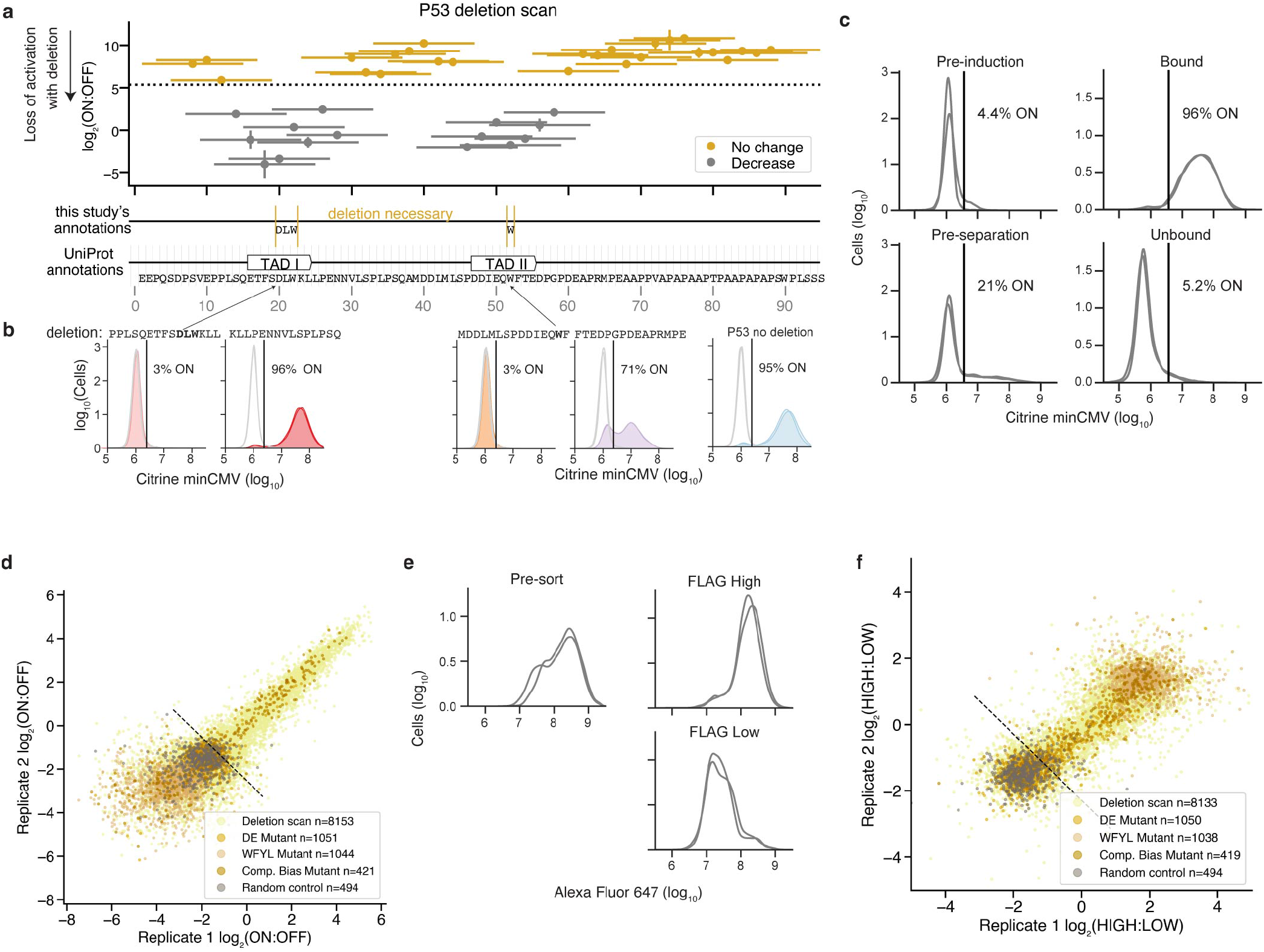
Mutant AD screen’s separation purity, reproducibility, and validation. **a**, Deletion scan across P53’s AD. If the deletion’s score is lower than 2 times the average standard error for measuring a deletion, it’s binned as “decrease”. Otherwise it’s binned as “no change.” **b**, Individual validations of 80 aa sequences including 15 aa deletions (deleted sequences shown above each panel). Untreated cells (gray) and dox-treated cells (colors) shown with two biological replicates in each condition. Vertical line is the citrine gate used to determine the fraction of cells ON (written above each distribution). **c**, Flow cytometry data showing citrine reporter distributions for the Mutant AD transcriptional activity screen on the day we induced localization with dox (Pre-induction), 2 days later on the day of magnetic separation (Pre-separation), and after separation using ProG DynaBeads that bind to the surface synthetic marker (Bound). Overlapping histograms are shown for 2 separately transduced biological replicates. The average percentage of cells ON is shown to the right of the vertical line showing the citrine level gate. A total of 1,000 ng/mL dox was added each day of dox treatment. **d**, Biological replicate Mutant AD transcriptional activity screen reproducibility. **e**, Alexa Fluor 647 distributions from anti-FLAG staining. **f**, Biological replicate Mutant AD protein expression screen reproducibility.

**Extended Data Fig. 6.**
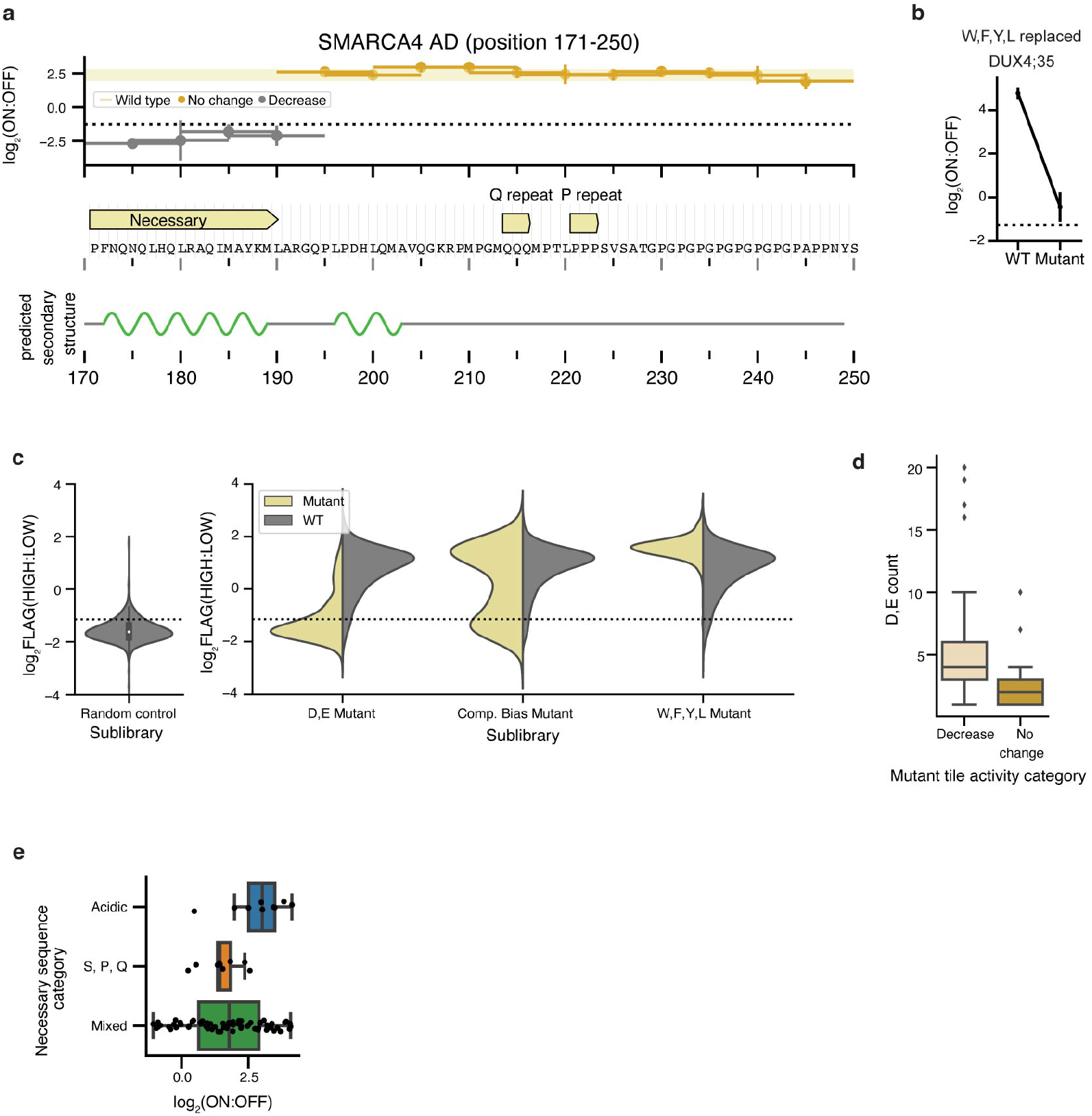
Mutant AD screen follow-up. **a**, Deletion scan across SMARCA4’s AD. AlphaFold’s predicted secondary structure (prediction from whole protein sequence) shown below where green regions are alpha helices. **b**, Line plot of average enrichment scores from two biological replicates. **c**, Violin plots of average FLAG enrichment scores from 2 biological replicates binned by each sublibrary. Dashed line represents the hit threshold. **d**, Boxplot of acidic count for each mutant’s activation category. Mann-Whitney one-sided U test, p-value=2.25e-3. **e**, Boxplot of average activation enrichment scores with IQR shown for tiles that contain a single necessary sequence across each category.

**Extended Data Fig. 7.**
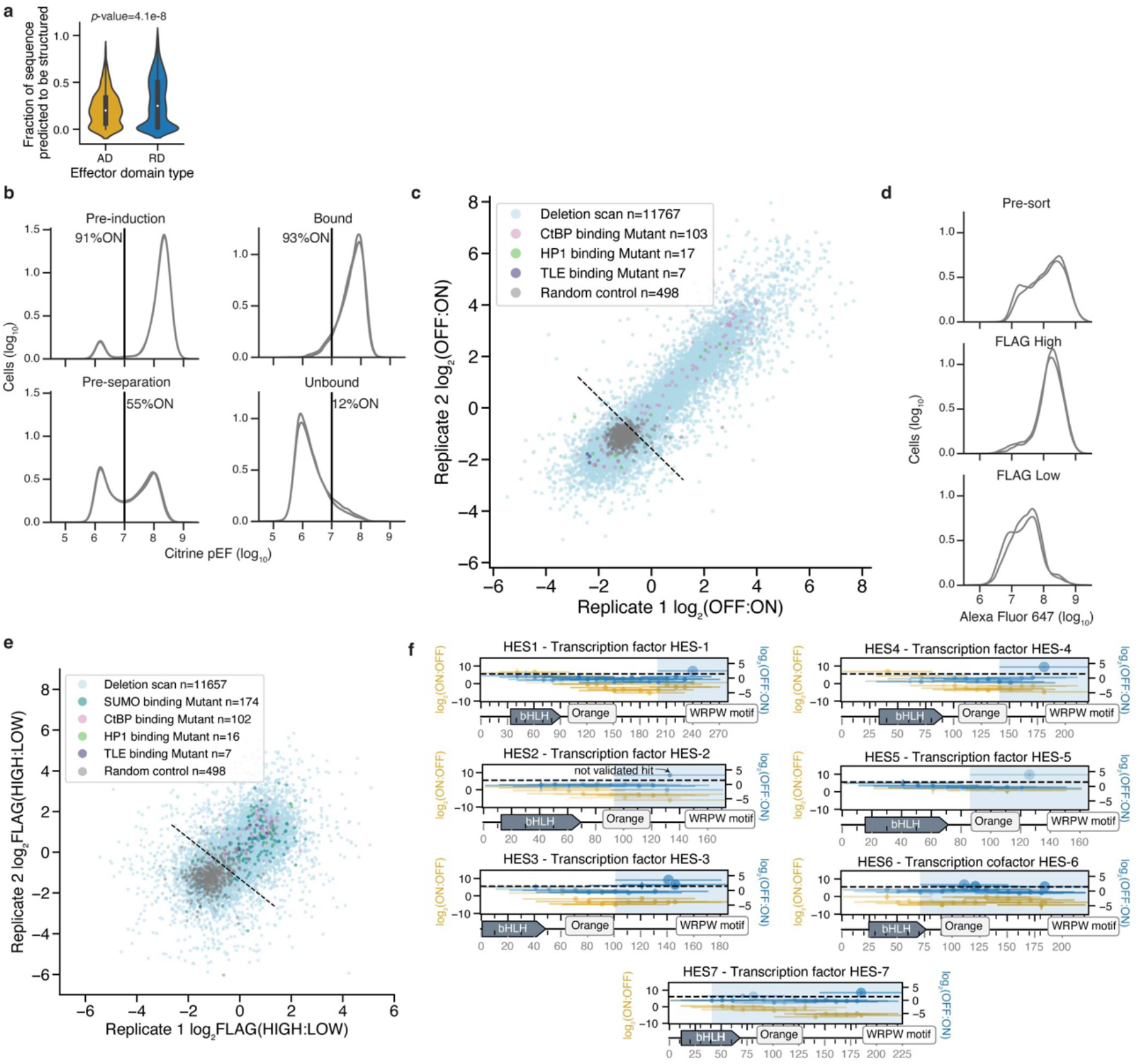
Distribution of tile’s predicted secondary structure, mutant RD screen’s separation purity and reproducibility, and HES family tiling plot examples. **a**, Distributions of activating and repressing tile’s fraction of sequence that’s predicted to be structured from AlphaFold’s predictions on the full length protein sequence. p-value=4.1e-8 (Mann Whitney U test, one-sided). **b**, Flow cytometry data showing citrine reporter distributions for the Mutant RD transcriptional activity screen on the day we induced localization with dox (Pre-induction), 5 days later on the day of magnetic separation (Pre-separation), and after separation using ProG DynaBeads that bind to the surface synthetic marker (Bound). Overlapping histograms are shown for 2 separately transduced biological replicates. The average percentage of cells ON is shown to the right of the vertical line showing the citrine level gate. A total of 1,000 ng/mL dox was added each day of dox treatment. **c**, Biological replicate Mutant RD transcriptional activity screen reproducibility. **d**, Alexa Fluor 647 staining distributions for the Mutant RD FLAG protein expression screen. **e**, Biological replicate Mutant RD protein expression screen reproducibility. **f**, Tiling plots for all 7 HES family members.

**Extended Data Fig. 8.**
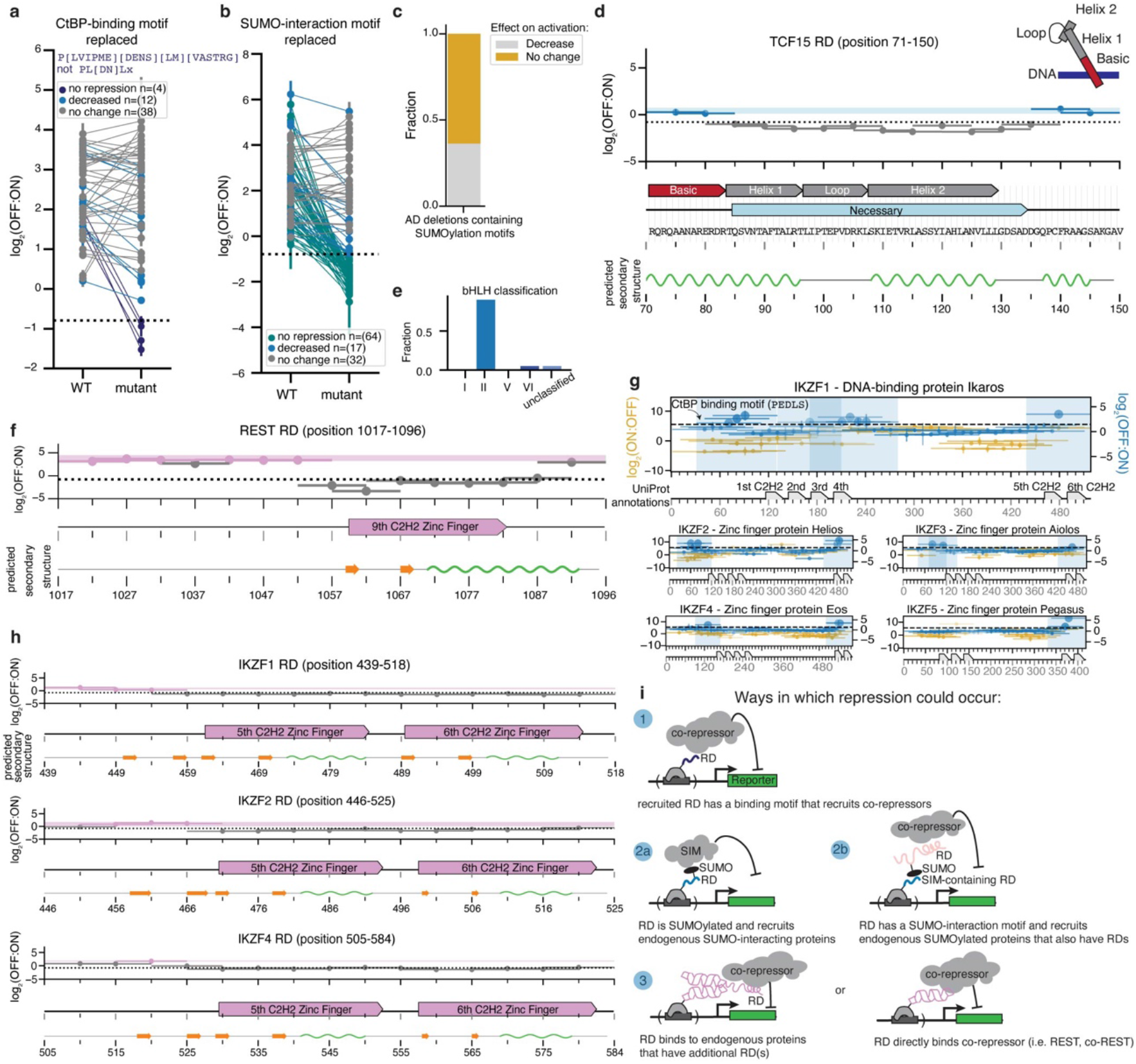
Mutant RD screen follow-up. **a**, Repression enrichment scores for a subset of repressing tiles that contain a relatively more flexible CtBP-binding motif (regex shown above), excluding the more refined CtBP-binding motif (regex shown on second line). Mutants have their binding motifs replaced with alanines. **b**, Repression enrichment scores for repressing tiles that contain a flexible SUMO-binding motif (fraction of non-hit sequences containing motif=0.155). 2 biological replicates shown with standard error bars. **c**, Fraction of AD deletion sequences containing a SUMOylation motif binned according to their effect on activity (yellow=no change on activation relative to WT, gray=decreased activation). 11 total ADs. **d**, Deletion scan across TCF15’s RD. AlphaFold’s predicted secondary structure (prediction from whole protein sequence) shown below where green regions are alpha helices. Annotations shown from protein accession NP_004600.3 **e**, Distribution of bHLH classifications of RDs overlapping bHLH UniProt annotations. Classifications taken from ref^33^. **f**, Deletion scan across REST’s RD. AlphaFold’s predicted secondary structure (prediction from whole protein sequence) shown below where green regions are alpha helices and orange arrows are beta sheets. **g**, Tiling plots for IKZF family members. **h**, Deletion scan across IKZF1, 2 and 4’s RDs. **i**, Cartoon model of potential mechanisms corresponding to the RD categories in Fig. 3f.

**Extended Data Fig. 9.**
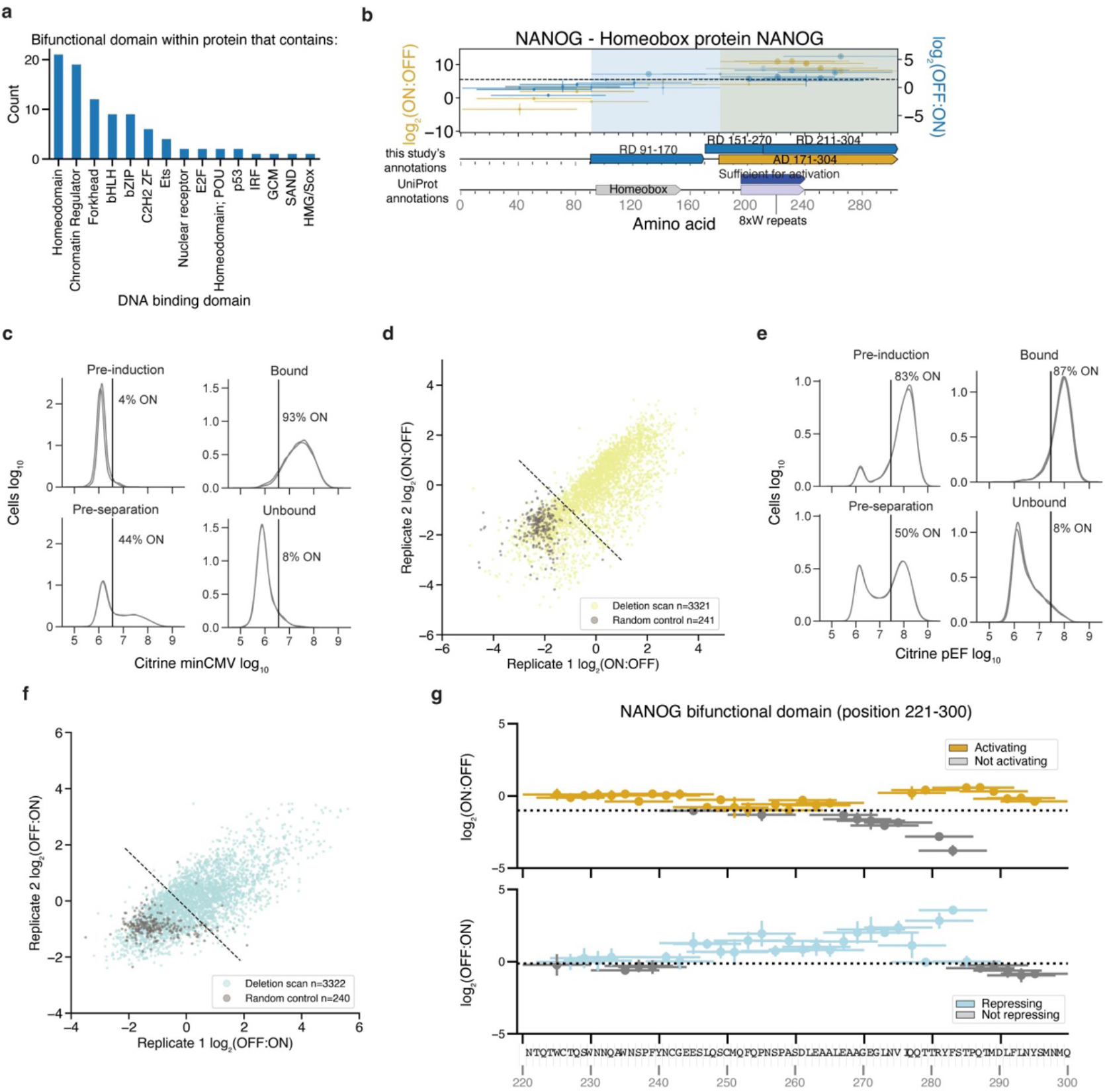
Bifunctional domain deletion scan screen’s separation purity, reproducibility, and examples. **a**, Counts of bifunctional domains from proteins that contain the following DNA binding domains. **b**, Tiling plot for NANOG. **c**, Flow cytometry data showing citrine reporter distributions for the bifunctional deletion scan minCMV promoter screen on the day we induced localization with dox (Pre-induction), 2 days later on the day of magnetic separation (Pre-separation), and after separation using ProG DynaBeads that bind to the surface synthetic marker (Bound). Overlapping histograms are shown for 2 separately transduced biological replicates. The average percentage of cells ON is shown to the right of the vertical line showing the citrine level gate. A total of 1,000 ng/mL dox was added each day of dox treatment. **d**, Biological replicate bifunctional deletion scan minCMV promoter screen reproducibility. **e**, Citrine reporter distributions for the bifunctional deletion scan pEF promoter screen, 5 days of induction (n=2). **f**, Biological replicate bifunctional deletion scan pEF promoter screen reproducibility. **g**, Example of a bifunctional domain from NANOG with independent activating and repressing regions (n=2). Note, deletion of the necessary sequence for activation, caused an increase in repression, and vice-versa.

**Extended Data Fig. 10.**
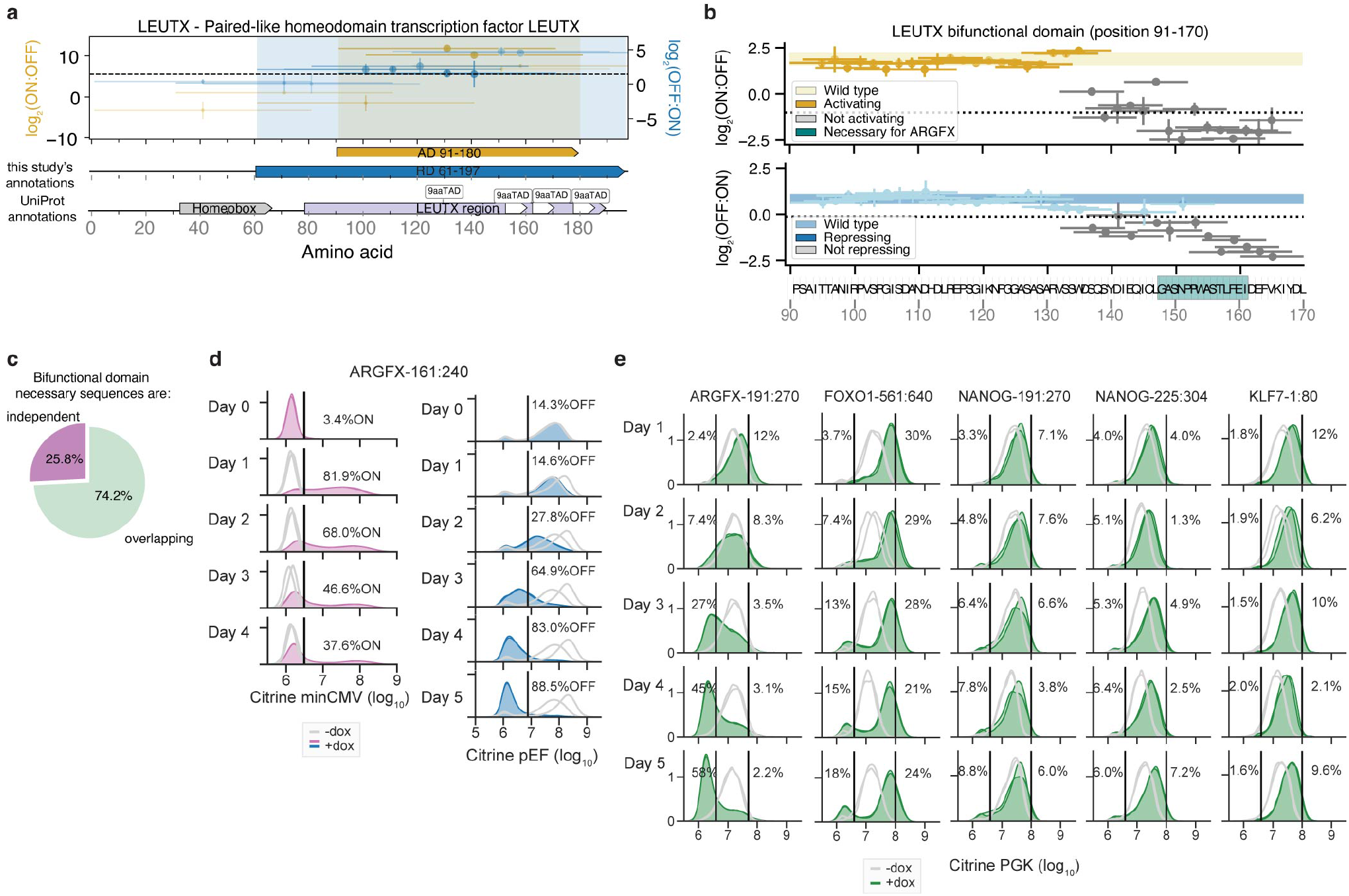
Examples of bifunctional domain sequences at three different promoters. **a**, Tiling plot for LEUTX. **b**, Deletion scan across one of LEUTX’s bifunctional tiles (n=2). The necessary sequence for gene family member, ARGFX, is highlighted in teal. **c**, Bifunctional domain necessary region location categories. Overlapping regions were defined as any tile that contained a deletion that was both necessary (below activity threshold) for activation and necessary for repression. **d**, Citrine distributions of ARGFX-161:240 recruited to minCMV (n=2, left), and recruited to pEF (n=2, right). **e**, Citrine distributions of bifunctional tiles identified from minCMV and pEF CRTF tiling screens recruited to PGK promoter (n=2).

## Supplementary Tables

**Supplementary Table 1: CRTF Tiles**

CRTF tiling library sequences and enrichment scores from the FLAG protein expression screen, minCMV, pEF, and PGK promoter screens, and the validation screens are attached in an Excel file.

**Supplementary Table 2: Domains from Tiles**

Activation and repression domain sequences and maximum tile enrichment scores are attached in an Excel file.

**Supplementary Table 3: Validations**

Individual validation flow cytometry data are attached in an Excel file.

**Supplementary Table 4: AD Mutants**

AD mutants library sequences and enrichment scores from the FLAG protein expression screen and minCMV promoter screen are attached in an Excel file.

**Supplementary Table 5: RD Mutants**

RD mutants library sequences and enrichment scores from the FLAG protein expression screen and pEF promoter screen are attached in an Excel file.

**Supplementary Table 6: Bifunctional Domains**

Bifunctional domains, deletion scan library sequences and enrichment scores from the FLAG protein expression screen, minCMV, and pEF promoter screens are attached in an Excel file.

## Methods

### Cell culture

All experiments presented here were carried out in K562 cells (ATCC, CCL-243, female). Cells were cultured in a controlled humidified incubator at 37C and 5% CO2, in RPMI 1640 (Gibco, 11-875-119) media supplemented with 10% FBS (Takara, 632180), and 1% Penicillin Streptomycin (Gibco, 15-140-122). HEK293T-LentiX (Takara Bio, 632180, female) cells, used to produce lentivirus, as described below, were grown in DMEM (Gibco, 10569069) media supplemented with 10% FBS (Takara, 632180) and 1% Penicillin Streptomycin Glutamine (Gibco, 10378016). minCMV and pEF reporter cell line generation is described in ref^4^. Briefly, pEF and minCMV promoter reporter cell lines were generated by TALEN-mediated homology-directed repair to integrate donor constructs (pEF promoter: Addgene #161927, minCMV promoter: Addgene #161928) into the *AAVS1* locus by electroporation of K562 cells with 1000 ng of reporter donor plasmid and 500 ng of each TALEN-L (Addgene #35431) and TALEN-R (Addgene #35432) plasmid (targeting upstream and downstream the intended DNA cleavage site, respectively). After 7 days, the cells were treated with 1000 ng/mL puromycin antibiotic for 5 days to select for a population where the donor was stably integrated in the intended locus. Fluorescent reporter expression was measured by microscopy and by flow cytometry. The PGK reporter cell line was generated by electroporation of K562 cells with 0.5 ug each of plasmids encoding the *AAVS1* TALENs and 1 ug of donor reporter plasmid using program T-016 on the Nucleofector 2b (Lonza, AAB-1001). Cells were treated with 0.5 ug/mL puromycin for one week to enrich for successful integrants. These cell lines were not authenticated. All cell lines tested negative for mycoplasma.

### TF tiling library design

1294 human transcription factors (TFs) were curated from ref^1^. We filtered out all KRAB-containing C2H2 zinc fingers, as our lab has previously screened the effector domains (KRAB, SCAN, DUF) of these proteins^4^. The canonical transcript of each gene was retrieved from Ensembl and chosen using the APPRIS principle transcript^53^. If no APPRIS tag was found, the transcript was chosen using the TSL principle transcript. If no TSL tag was found, the longest transcript with a protein coding CDS was retrieved. The coding sequences were divided into 80 amino acid (aa) tiles with a 10 aa sliding window. For each gene, a final tile was included spanning from 80 aa upstream of the last residue to that last residue, such that the C-terminal region would be included in the library. Duplicate sequences were removed, sequences were codon matched for human codon usage, 7xC homopolymers were removed, BsmBI restriction sites were removed, rare codons (less than 10% frequency) were avoided, and the GC content was constrained to be between 20% and 75% in every 50 nucleotide window (performed with DNA chisel^54^). To improve the coverage of this large library, we subdivided into 3 smaller sub-libraries based on the three major classes of TFs: a 25,032 C2H2 ZF sub-library including all 406 C2H2 ZF TFs, a 9,757 Homeodomain and bHLH sub-library including all 304 Homeodomain and bHLH TFs, and a 31,664 member sub-library containing the rest of the 583 TFs.

1000 random controls of 80 amino acids lacking stop codons were computationally generated as controls using the DNA chisel package’s *random_dna_sequence* function and included in each sub-library. 473 sequences that were found to be non-activators and 42 sequences that were found to be activators in our lab’s previous minCMV Nuclear Pfam screen^4^ were included as negative and positive controls. We made use of alternative codon usage *(match_codon_usage*, and *use_best_codon* functions) to re-code the controls in each sub-library in order to give ourselves the option of pooling the 3 sub-libraries and running the library as one 73,288 element screen.

100 additional controls were added to each sub-library to serve as fiduciary markers to aid comparing separately run screens. These controls were not recoded in each sub-library, and thus were repeated when pooling sub-libraries.

50 activation domains from 45 proteins involved in transcriptional activation were curated from UniProt^3^. We queried the UniProt database for human proteins whose regions, motifs or annotations included the term “transcriptional activation.” We then filtered for ADs that ranged in length from 30 to 95 aa. For ADs shorter than 95 aa, we extended the protein sequence equally on either side until it reached 95 aa. The protein sequences were reverse translated and further divided into 95 aa sequences with 15 aa deletions positioned with a 2 aa sliding window. Duplicate sequences were removed, sequences were codon matched for human codon usage, 7xC homopolymers were removed, BsmBI restriction sites were removed, rare codons (less than 10% frequency) were avoided, and the GC content was constrained to be between 20% and 75% in every 50 nucleotide window (performed with DNA chisel^54^. 50 yeast Gcn4 controls were added, which included previously studied deletions^29^. 2,024 library elements in total were added to the 31,664 element TF tiling sub-library.

### CR tiling library design

Candidate genes were initially chosen by including all members of the EpiFactors database, genes with gene name prefixes that matched any genes in the EpiFactors database, and genes with any of the following GO terms: GO:000785 (chromatin), GO:0035561 (regulation of chromatin binding), GO:0016569 (covalent chromatin modification), GO:1902275 (regulation of chromatin organization), GO:0003682 (chromatin binding), GO:0042393 (histone binding), GO:0016570 (histone modification), and GO:0006304 (DNA modification). Genes present in prior Silencer tiling screens^4^ and genes present in the present TF tiling screen were then filtered out. Biomart was used to identify and retrieve the canonical transcript, and chosen by (in order of priority) the APPRIS principal transcript, the TSL principal transcript, or the longest transcript with a protein coding CDS. Tiles for each of these DNA sequences were generated using the same 80 amino acid tile/10 amino acid sliding window approach as the TF tiling library. Duplicate sequences were removed, DNA hairpins and 7xC homopolymers were removed, and sequences were codon matched for human codon usage with GC content being constrained to be between 20% and 75% globally and between 25% and 65% in any 50-bp window. In order to improve the coverage while performing the screen, this 51,297 element library was split into two sub-libraries: a 38,241 element CR Tiling Main sub-library and an 13,056 element CR Tiling Extended sub-library. Computationally generated random negative controls, negative control tiles from the DMD protein screened in prior Nuclear Pfam screens^4^, and fiduciary marker controls were added to each sub-library: 1,700 elements to the Main sub-library and 3,700 elements to the Extended sub-library. These controls were not re-coded, and thus were repeated when pooling sub-libraries.

### Library filtering

Since we pooled the sub-libraries and screened them as one large pool, several of the control sub-libraries, that were not re-coded, wound up being repeated in the pool several times. We noticed sequences that were repeated fewer times had enrichment scores closer to what was observed previously. But sequences that were repeated upwards of five times had systematically lower enrichment scores than what was expected from previous screens, likely due to PCR bias. We removed all repeated control elements and instead relied on individual validations to confirm our screens worked. Additionally, there was a computational error in removing BsmBI sites from the CR tiling library, resulting in some sequences having accidental restriction cut sites in the middle of the ORF. We removed these sequences from further analysis and supplementary tables.

### Activating hits validation library design

1,055 putative hit tiles were chosen by selecting all tiles where both biological replicates were recovered and had activation enrichment scores above 5.365 (determined by 2 standard deviations above random controls). We included 200 randomly selected random negative controls that were poorly expressed (expression threshold = -1.427) and 100 randomly selected non-hit tiles that had no activity in both the minCMV and the pEF CRTF tiling screens. There were 1,355 total library elements.

### Repressing hits validation library design

9,438 putative hit tiles were chosen by selecting all tiles where both biological replicates were recovered and had pEF repression enrichment scores above 1.433 or had a PGK repression enrichment scores above 0.880 (determined from 3 standard deviations above random controls). We included 500 randomly selected random negative controls that were poorly expressed (expression threshold = -1.427) and 100 randomly selected non-hit tiles that had no activity in the minCMV, pEF nor PGK CRTF tiling screens. There were 10,038 total library elements.

### AD mutants library design

We defined compositional bias as any residue that represented more than 15% of the sequence (more than 12 residues). We took 424 compositionally biased tiles and replaced all residues with alanine. We took 1055 aromatic or leucine-containing tiles and replaced all Ws, Fs, Ys, and Ls with alanine. We took 1052 acidic residue-containing tiles and replaced all Ds and Es with alanine. 51 tiles that contained the “LxxLL” motif (ELM accession: ELME000045, regex pattern = [^P]L[^P][^P]LL[^P]) we replaced with alanine. 22 tiles that contained the “WW” motif (ELM accession: ELME000003, regex pattern = PP.Y) we replaced with alanine. 8205 deletions were designed by systematically removing 10 aa chunks, with a sliding window of 5 aa from 547 max activating tiles. All mutated sequences were reverse translated into DNA sequences using a probabilistic codon optimization algorithm, such that each DNA sequence contains some variation beyond the substituted residues, which improves the ability to unambiguously align sequencing reads to unique library members. The 1055 putative hit tiles were included as positive controls (slightly more activating tiles than we report in the main text because these libraries were designed before we screened the validation library). We included 500 randomly selected random negative controls that were poorly expressed (expression threshold = -1.427). There were 12,364 total library elements.

### RD mutants library design

12,000 deletions were designed by systematically removing 10 aa chunks, with a sliding window of 5 aa of the maximum tile from 800 putative RDs that were hits in both PGK and pEF CRTF tiling screens (slightly more RDs than we report in the main text because these libraries were designed before we screened the validation library). All mutated sequences were reverse translated into DNA using the method described above. The 1,593 putative hit tiles were included as positive controls. We took 644 compositionally biased tiles and replaced all residues with alanine. We replaced with alanines all the following motifs: 104 CtBP interaction motif containing tiles (ELM accession: ELME0000098); 18 HP1 interaction motif containing tiles (ELM accession: ELME000141); 9 “ARKS” motif containing tiles (ELM accession: DRAFT - LIG_CHROMO); 180 SUMO interaction motif containing tiles (ELM accession: ELME000335); and 7 WRPW motif containing tiles (ELM accession: ELME000104). We included 500 randomly selected random negative controls that were poorly expressed (expression threshold = -1.427). There were 15,055 total library elements.

### Bifunctional deletion scan library design

3,331 deletions were created by systematically removing 10 aa chunks, with a sliding window of 2 aa from 96 bifunctional activating and repressing tiles. All mutated sequences were reverse translated into DNA sequences using a method described above. We included the WT bifunctional tiles and 250 randomly selected random negative controls that were poorly expressed (expression threshold = -1.427). There were 3,674 total library elements.

### Library cloning

Oligonucleotides with lengths up to 300 nucleotides were synthesized as pooled libraries (Twist Biosciences) and then PCR amplified. 6x 50 ul reactions were set up in a clean PCR hood to avoid amplifying contaminating DNA. For each reaction, we used either 5 or 10 ng of template, 1 ul of each 10 mM primer, 1 ul of Herculase II polymerase (Agilent), 1 ul of DMSO, 1 ul of 10 mM dNTPs, and 10 ul of 5x Herculase buffer. The thermocycling protocol was 3 minutes at 98C, then cycles of 98C for 20 s, 61C for 20 s, 72C for 30 s, and then a final step of 72C for 3 minutes. The default cycle number was 20x, and this was optimized for each library to find the lowest cycle that resulted in a clean visible product for gel extraction (in practice, 23 cycles was the maximum when small libraries were represented in large pools). After PCR, the resulting dsDNA libraries were gel extracted by loading a 2% TAE gel, excising the band at the expected length (around 300 bp), and using a QIAgen gel extraction kit. The libraries were cloned into a lentiviral recruitment vector pJT126 (Addgene #161926) with 4-16x 10 ul Golden-Gate reactions (75 ng of pre-digested and gel-extracted backbone plasmid, 5 ng of library (2:1 molar ratio of insert:backbone), 2uL of 10x T4 Ligase Buffer, and 1uL of NEB Golden Gate Assembly Kit (BsmBI-V2)) with 65 cycles of digestion at 42C and ligation at 16C for 5 minutes each, followed by a final 5 minute digestion at 42C and then 20 minutes of heat inactivation at 70C. The reactions were then pooled and purified with MinElute columns (QIAgen), eluting in 6 ul of ddH2O. 2 ul per tube was transformed into two tubes of 50 ml of Endura electrocompetent cells (Lucigen, Cat#60242-2) following the manufacturer’s instructions. After recovery, the cells were plated on 1-8 large 10’’x10’’ LB plates with carbenicillin. After overnight growth in a warm room, the bacterial colonies were scraped into a collection bottle and plasmid pools were extracted with a Hi-Speed Plasmid Maxiprep kit (QIAgen). 2-3 small plates were prepared in parallel with diluted transformed cells in order to count colonies and confirm the transformation efficiency was sufficient to maintain at least 20x library coverage. To determine the quality of the libraries, the putative effector domains were amplified from the plasmid pool by PCR with primers with extensions that include Illumina adapters and sequenced. The PCR and sequencing protocol were the same as described below for sequencing from genomic DNA, except these PCRs use 10 ng of input DNA and 17 cycles. These sequencing datasets were analyzed as described below to determine the uniformity of coverage and synthesis quality of the libraries. In addition, 20-30 colonies from the transformations were Sanger sequenced (Quintara) to estimate the cloning efficiency and the proportion of empty backbone plasmids in the pools.

### Pooled delivery of library in human cells using lentivirus

Large scale lentivirus production and spinfection of K562 cells were performed as follows: To generate sufficient lentivirus to infect the libraries into K562 cells, we plated HEK293T cells on 1-12 15-cm tissue culture plates. On each plate, 8.8 × 10^6^ HEK293T cells were plated in 30 mL of DMEM, grown overnight, and then transfected with 8 ug of an equimolar mixture of the three third-generation packaging plasmids (pMD2.G, psPAX2, pMDLg/pRRE) and 8 ug of rTetR-domain library vectors using 50 mL of polyethylenimine (PEI, Polysciences #23966). pMD2.G (Addgene plasmid #12259; http://addgene.org/12259), psPAX2 (Addgene plasmid #12260; http://addgene.org/12260), and pMDLg/pRRE (Addgene plasmid #12251; http://addgene.org/12251) were gifts from Didier Trono. After 48 hours and 72 hours of incubation, lentivirus was harvested. We filtered the pooled lentivirus through a 0.45-mm PVDF filter (Millipore) to remove any cellular debris. K562 reporter cells were infected with the lentiviral library by spinfection for 2 hours, with two separate biological replicates infected. Infected cells grew for 2 days and then the cells were selected with blasticidin (10 mg/mL, Gibco). Infection and selection efficiency were monitored each day using flow cytometry to measure mCherry (Biorad ZE5). Cells were maintained in spinner flasks in log growth conditions each day by diluting cell concentrations back to a 5 × 10^5^ cells/mL. We aimed for 600x infection coverage and our lowest infection coverage was 130x. We aimed to have 2-10,000x maintenance coverage. On day 8 post-infection, recruitment was induced by treating the cells with 1000 ng/ml doxycycline (Fisher Scientific) for either 2 days for activation or 5 days for repression.

### Magnetic separation

At each time point, cells were spun down at 300 x *g* for 5 minutes and media was aspirated. Cells were then resuspended in the same volume of PBS (GIBCO) and the spin down and aspiration was repeated, to wash the cells and remove any IgG from serum. Dynabeads M-280 Protein G (ThermoFisher, 10003D) were resuspended by vortexing for 30 s. 50 mL of blocking buffer was prepared per 2 × 10^8^ cells by adding 1 g of biotin-free BSA (Sigma Aldrich) and 200 mL of 0.5 M pH 8.0 EDTA into DPBS (GIBCO), vacuum filtering with a 0.22-mm filter (Millipore), and then kept on ice. For all activation screens, 30 uL of beads was prepared for every 1 × 10^7^ cells, 60 uL of beads/10 million cells for the pEF CRTF tiles, PGK CRTF tiles, and minCMV bifunctional deletion scan screens, 120 uL of beads/10 million cells for the pEF validation, 90 uL of beads/10 million cells for the RD Mutants and pEF bifunctional deletion scan screens. Magnetic separation was performed as previously described in ref^4^.

### FLAG staining for protein expression

The expression level measurements for the CRTF tiling library were made in K562 minCMV cells (with citrine OFF). 4 × 10^8^ cells per biological replicate were used after 7 days of blasticidin selection (10 mg/mL, Gibco), which was 9 days post-infection. 4 × 10^7^ control K562-JT039 cells (citrine ON, no lentiviral infection) were spiked into each replicate. Fix Buffer I (BD Biosciences, BDB557870) was preheated to 37C for 15 minutes and Permeabilization Buffer III (BD Biosciences, BDB558050) and PBS (GIBCO) with 10% FBS (Hyclone) were chilled on ice. The library of cells expressing domains was collected and cell density was counted by flow cytometry (Biorad ZE5). To fix, cells were resuspended in a volume of Fix Buffer I (BD Biosciences, BDB557870) corresponding to pellet volume, with 20 mL per 1 million cells, at 37C for 10 - 15 minutes. Cells were washed with 1 mL of cold PBS containing 10% FBS, spun down at 500 3 g for 5 minutes and then supernatant was aspirated. Cells were permeabilized for 30 minutes on ice using cold BD Permeabilization Buffer III (BD Biosciences, BDB558050), with 20 mL per 1 million cells, which was added slowly and mixed by vortexing. Cells were then washed twice in 1 mL PBS+10% FBS, as before, and then supernatant was aspirated. Antibody staining was performed for 1 hour at room temperature, protected from light, using 5 uL / 1 × 10^6^ cells of a-FLAG-Alexa647 (RNDsystems, IC8529R). We then washed the cells and resuspended them at a concentration of 3 × 10^7^ cells / ml in PBS+10%FBS. Cells were sorted into two bins based on the level of APC-A and mCherry fluorescence (Sony SH800S) after gating for viable cells. A small number of unstained control cells was also analyzed on the sorter to confirm staining was above background. The spike-in citrine positive cells were used to measure the background level of staining in cells known to lack the 3XFLAG tag, and the gate for sorting was drawn above that level. After sorting, the cellular coverage was ∼2000x. The sorted cells were spun down at 500 x g for 5 minutes and then resuspended in PBS. Genomic DNA extraction was performed following the manufacturer’s instructions (QIAgen Blood Midi kit was used for samples with > 1 × 10^7^ cells) with one modification: the Proteinase K + AL buffer incubation was performed overnight at 56C.

### Library preparation and sequencing

Genomic DNA was extracted with the QIAgen Blood Maxi Kit following the manufacturer’s instructions with up to 1 × 10^8^ cells per column. DNA was eluted in EB and not AE to avoid subsequent PCR inhibition. The domain sequences were amplified by PCR with primers containing Illumina adapters as extensions. A test PCR was performed using 5 ug of genomic DNA in a 50 mL (half-size) reaction to verify if the PCR conditions would result in a visible band at the expected size for each sample. Then, 3 - 48x 100 uL reactions were set up on ice (in a clean PCR hood to avoid amplifying contaminating DNA), with the number of reactions depending on the amount of genomic DNA available in each experiment. 10 ug of genomic DNA, 0.5 mL of each 100 mM primer, and 50 mL of NEBnext Ultra 2x Master Mix (NEB) was used in each reaction. The thermocycling protocol was to preheat the thermocycler to 98C, then add samples for 3 minutes at 98C, then an optimized number of cycles of 98C for 10 s, 63C for 30 s, 72C for 30 s, and then a final step of 72C for 2 minutes. All subsequent steps were performed outside the PCR hood. The PCR reactions were pooled and 145 uL were run on a 2% TAE gel, the library band around 395 bp was cut out, and DNA was purified using the QIAquick Gel Extraction kit (QIAgen) with a 30 ul elution into non-stick tubes (Ambion). A confirmatory gel was run to verify that small products were removed. These libraries were then quantified with a Qubit HS kit (Thermo Fisher) and sequenced on an Illumina HiSeq (2×150).

### Computing enrichments and hits thresholds

Sequencing reads were demultiplexed using bcl2fastq (Illumina). A Bowtie reference was generated using the designed library sequences with the script ‘makeIndices.py’ (HT-Recruit Analyze package) and reads were aligned with 0 mismatch allowance using the script ‘makeCounts.py’. The enrichments for each domain between OFF and ON (or FLAGhigh and FLAGlow) samples were computed using the script ‘makeRhos.py’. Domains with < 5 reads in both samples for a given replicate were dropped from that replicate (assigned 0 counts), whereas domains with < 5 reads in one sample would have those reads adjusted to 5 in order to avoid the inflation of enrichment values from low depth.

For all of the screens, domains with < 20 counts in both conditions of a given replicate were filtered out of downstream analysis. For the expression screens, well-expressed tiles were those with a log2(FLAGhigh:FLAGlow) 1 standard deviation above the median of the random controls. For the CRTF tiling repressor screens, hits were tiles with enrichment scores 3 standard deviations above the mean of the poorly expressed random controls. For the minCMV CRTF tiling, pEF Bifunctional deletion scan, and minCMV bifunctional deletion scan screens, hits were proteins with enrichment scores 2 standard deviations above the mean of the poorly expressed random controls. For the validation and mutant screens, hits were proteins with enrichment scores 1 standard deviation above the mean of the poorly expressed random controls.

### Annotation of domains from tiles

Tiles must have been hits in both the CRTF tiling and validation screens in order to have been considered potential effector domains. A domain started anywhere the previous tile was not a hit. If the previous tile was not a hit because it was not expressed, and if the antepenultimate (previous, previous) tile was a hit, then that tile was not considered the start, and it was recovered into the middle of the domain. A domain ended anywhere the next successive tile was not a hit. If the next tile was not a hit because it was not expressed, and the following tile was a hit, then that tile was not considered the end, and it was recovered into the middle of the domain. Domains started at the first residue of the first tile and extended until the last residue of the last tile within the domain.

### Individual recruitment assays

Protein fragments were cloned as a fusion with rTetR upstream of a T2A-mCherry-BSD marker, using GoldenGate cloning in the backbone pJT126 (Addgene #161926). K562 citrine reporter cells were then transduced with each lentiviral vector and, 3 days later, selected with blasticidin (10 mg/mL) until > 80% of the cells were mCherry positive (6-9 days). Cells were split into separate wells of a 24-well plate and either treated with doxycycline (Fisher Scientific) or left untreated. Time points were measured by flow cytometry analysis of >10,000 cells (Biorad ZE5). Doxycycline was assumed to be degraded each day, so fresh doxycycline media was added each day of the timecourse.

### Western blots

5-10 million cells were lysed in lysis buffer (1% Triton X-100, 150mM NaCl, 50mM Tris pH 7.5, Protease inhibitor cocktail). Protein amounts were quantified using the Pierce BCA Protein Assay kit (Bio-Rad). Equal amounts were loaded onto a gel and transferred to a PVDF membrane. Membrane was probed using FLAG M2 monoclonal antibody (1:1000, mouse, Sigma-Aldrich, F1804) and Histone 3 antibody (1:1000, mouse, Abcam, AB1791) as primary antibodies. Goat anti-mouse IRDye 680 RD and goat anti-rabbit IRDye 800CW (1:20,000 dilution, LICOR Biosciences, cat nos. 926-68070 and 926-32211, respectively) were used as secondary antibodies. Blots were imaged on an iBright (Thermo Scientific). Band intensities were quantified using ImageJ.

### Data analysis and statistics

All statistical analyses and graphical displays were performed in Python^55^ (v. 3.8.5). Enrichment scores shown in all figures (aside from replicate plots) are the average across two separately transduced biological replicates. The p-values, statistical tests used, and n are indicated in the figure legends.

### Flow cytometry analysis

Data were analyzed using Cytoflow (https://github.com/bpteague/cytoflow) and custom Python scripts. Events were gated for viability and mCherry as a delivery marker. To compute a fraction of ON cells during doxycycline treatment, we fit a Gaussian model to the untreated rTetR-only negative control cells which fits the OFF peak, and then set a threshold that was 2 standard deviations above the mean of the OFF peak in order to label cells that have activated as ON. We do the same for computing the fraction of OFF cells in repressor validations but fit a two component Gaussian and set a threshold that was 2 standard deviations below the mean of the ON peak. A logistic model, including a scale parameter, was fit to the validation and screen data using SciPy’s curve fit function.

## Data availability

All raw NGS data and associated processed data generated in this study will be deposited in the NCBI GEO database upon publication.

## Code availability

The HT-recruit Analyze software for processing high-throughput recruitment assay and high-throughput protein expression assays are available on GitHub (https://github.com/bintulab/HT-recruit-Analyze).

All custom codes used for data processing and computational analyses are available from the authors upon request.

## Biological materials availability

Oligonucleotide libraries are available upon request.

## Acknowledgements

We thank Michaela Hinks and members of our laboratories for helpful conversations and assistance. This work was supported by NIH-NIGMS R35M128947 (L.B.), NSF GRFP DGE-1656518 (N.D.), NIH-NIDDK F99/K00 F99DK126120 (J.T.), Stanford Bio-X Bowes Fellowship (P.S.), Stanford School of Medicine Dean’s Fund, (C.A.), NIH-NIGMS 5T32GM007365-45 (A.M.), Stanford Interdisciplinary Graduate Fellowship affiliated with Stanford Bio-X (A.M.), NIH Director’s New Innovator Award (1DP2HD08406901) (M.B.), and the BWF-CASI Award (L.B.)

## Authorship contributions

N.D. and L.B. designed the study, with significant intellectual contributions from P.S. and A.M. P.S and N.D designed the TF tiling libraries, A.M designed the CR tiling libraries, both with contributions from J.T, M.C.B. and L.B. N.D. designed all other libraries with contributions from J.T, A.M, P.S, M.C.B. and L.B. N.D. screened the CRTF minCMV and FLAG libraries with assistance from P.S and J.T., Aradhana, K.S screened the CRTF pEF and PGK promoter libraries. N.D. performed all other screens. N.D. analyzed the data, with assistance from L.B. I.L., C.A., and N.D. performed individual recruitment assay experiments. N.D. performed Western blot experiments. C.L. generated the PGK cell line. N.D. and L.B. wrote the manuscript, with significant contributions from J.T. and C.L, along with contributions from all authors. P.F, M.C.B. and L.B. supervised the project.

